# Cell-Autonomous and Systemic Circadian Regulation of Gene Expression in Adipocytes

**DOI:** 10.64898/2026.01.24.701487

**Authors:** Jennifer M. Worthen, Armina-Lyn M. Frederick, Phillip A. Dumesic, Jennifer J. Loros, Jay C. Dunlap

## Abstract

Circadian clocks strongly influence adipocyte biology. Studying adipocyte circadian regulation in the absence of organismal cues isolates cell-autonomous clock control, providing insight into mechanisms relevant to metabolic disease. To resolve circadian core biological programs, we deeply sequenced bulk RNA from inguinal-derived, *in vitro*-differentiated adipocytes (IVDAs) over 2.5 days and compared clock-controlled genes (CCGs) with existing mouse supraclavicular brown (BAT) and epididymal white (eWAT) adipose tissue circadian time-course datasets. Using Phase Set Enrichment Analysis (PSEA), we report 21% of the protein-coding transcriptome is rhythmic in IVDAs. Intrinsic circadian regulation governs key processes in energy metabolism, molecular transport and transcription. Integration with *in vivo* datasets reveals that BAT and IVDAs exhibit more cohesive rhythmic pathways than does eWAT, clustering around the late-night early-morning transition. To explore how these pathways may be regulated, we reexamined a recent interscapular BAT cistrome dataset. Using IVDA transcription factors that were phase-aligned (≤4hr) with *in vivo* as input, motif enrichment analysis revealed two temporally distinct regulatory programs; an early E-box activator ARNT-family/bHLH-PAS program was enriched for transcriptional regulation, RNA metabolism, and signaling pathways, and a late nuclear receptor-associated program enriched for energy metabolism, phospholipid biosynthesis, mitochondrial function, ECM organization, and nuclear receptor signaling. Overall, we identify novel rhythmic transcripts and define cell-autonomous circadian programs in adipocytes whose timing is further sculpted by systemic cues *in vivo*. Because obesity is associated with adipocyte hypertrophy and hyperplasia, processes likely influenced by circadian regulation, these findings advance our understanding of clock-controlled adipocyte metabolism and its contribution to metabolic dysfunction.

## INTRODUCTION

The circadian clock plays a central role in regulating metabolism by synchronizing physiological processes to the 24h daily cycle (Bass, 2012; Hurley et al., 2016; Schrader et al., 2024). In nearly every animal cell, this internal timekeeper coordinates key functions such as energy expenditure and intermediary metabolism (van Rosmalen et al., 2024; Neufeld-Cohen et al., 2016; Malik et al., 2024). While studies of adipose tissue *in vivo* have established the rhythmic nature of adipose tissue function, the extent to which these rhythms are cell-intrinsic versus systemically driven remains unclear. Adipocytes integrate nutrient availability, hormonal signals, and energetic demand and retain the capacity for autonomous circadian regulation. Clocks in adipose tissues drive rhythmic expression of genes and proteins critical to nutrient metabolism, signaling, and secretion, which influence glucose homeostasis and energy balance (Zvonic et al., 2006; Rosen and Spiegelman, 2006; Hepler and Bass, 2023). Adipose tissue is composed of several cell types, including mesenchymal stem cells, immune cells and vascular cells, with distinct anatomical depots serving specialized functions (Sakers et al., 2022; Yang Loureiro et al., 2022; Corvera et al., 2026). In addition, recent single-cell and single-nucleus studies have demonstrated substantial heterogeneity among adipocytes within individual depots, revealing multiple transcriptional states and functional subpopulations (Loft et al., 2025). While all white adipose tissue (WAT) stores energy as triglycerides in lipid droplets, brown adipose tissue (BAT) is also dense with mitochondria and constitutively capable of dissipating chemical energy as heat by adaptive non-shivering thermogenesis (ANST) (Bunk et al., 2025; Rahbani et al., 2024; Sun et al., 2021; Cannon and Nedergaard, 2004; Puigserver et al., 1998). Interspersed throughout inguinal WAT (iWAT) and other depots is a subtype transiently capable of ANST, referred to as “brown in white” or beige (Vargas-Castillo et al., 2024; Bertholet et al., 2017; reviewed in Inagaki et al., 2016). Over-feeding or cold exposure via sympathetic nervous system activation induces a metabolic sink via H^+^ leakage across the mitochondrial membrane or futile phosphorylation cycling, and this is retained through epigenomic memory (Roh et al., 2018). While the physiological relevance and mechanisms underlying this plasticity are debated (Cypess et al., 2025), the upregulation of an ANST-like response in adipocytes is, in part, driven by cell-autonomous sensing of ambient cold (Ye et al., 2013).

The circadian oscillator comprises a self-regulating transcriptional-translational feedback loop (TTFL). The core clock proteins CLOCK and BMAL1 form a heterodimer that enters the nucleus and binds to E-box sequences on target clock-controlled genes (CCGs) to initiate transcription (Takahashi, 2017; Van Drunen and Eckel-Mahan, 2021; Rasmussen et al., 2022). Upregulated genes include paralogs of Cryptochrome (*Cry*) and Period (*Per*), whose products form multimeric complexes with casein kinase 1d/e; these interact with CLOCK:BMAL1, resulting in their inhibition by blocking the activity of CLOCK and BMAL1 at E-boxes and thereby also reducing *Per/Cry* expression (Smyllie et al., 2025). Serial phosphorylation of the CRY and PER proteins by CK1 eventually relieves inhibition, allowing the cycle to reinitiate *Cry* and *Per* expression (Cao et al., 2021; Narasimamurthy and Virshup, 2021; Philpott et al., 2023; Ricci et al., 2025).

Disruption of circadian rhythms is strongly associated with metabolic dysfunction, including obesity and insulin resistance. When food or physical activity, very strong temporal cues, are shifted out of the active phase and into the resting phase, body composition as well as other parameters of metabolic health are altered (Acosta-Rodríguez et al., 2022; Acosta-Rodríguez et al., 2024), yet it remains unclear which aspects of adipocyte metabolic regulation arise from intrinsic clock control versus systemic cues including feeding, temperature, and neuroendocrine input (Allada and Bass, 2021; Fekry and Eckel-Mahan, 2022, Schrader et al., 2024). Resolving this distinction is essential for understanding how temporal regulation of adipocyte metabolism contributes to health and disease (Deota et al., 2025). Indeed, circadian regulation in adipose tissue has significant implications. In mice, chronic circadian shifts drive WAT hypertrophy, inflammation, and disrupted metabolic regulation, resulting in obesity-like transcriptional changes and impaired insulin signaling (Xiong et al., 2021). Circadian disruptors like shift work and jet lag alter adipokine expression and secretion, linking disrupted rhythms to metabolic dysfunction (Kettner et al., 2015; Scheer et al., 2009; Kiehn et al., 2017; Samanta and Ali, 2022). Brown adipose-specific deletion of *Arntl/Bmal1* disrupts thermogenesis and energy expenditure and increases susceptibility to developing obesity when mice are fed a high fat diet (Hasan et al., 2021). Characterizing the circadian regulation of target pathways offers insight into the pathogenesis and treatment of metabolic syndrome.

Here, we isolate adipocyte circadian biology from organismal context by comparing circadian transcriptomes of *in vitro* differentiated adipocytes (IVDAs) with *in vivo* brown and white adipose depots. This study used developmentally equivalent cells but excludes input from other cell types and from systemic influences such as feeding, hormones, or body temperature. Accordingly, our experimental design is optimized to capture cell-intrinsic circadian programs in differentiated adipocytes. Our differentiation protocol pushes cells into a committed adipocyte state and does not attempt to capture the adipocyte subtype heterogeneity revealed by single-cell profiling *in vivo* (Maniyadath et al., 2023). Through the systematic comparison of our deeply sequenced transcriptome with comparable datasets derived from whole animals (Zhang et al., 2014), we find that the cell-intrinsic circadian clock regulates the gene expression of fuel oxidation, adipose remodeling, and intra-organ communication programs. We have focused on identifying which pathways rely on systemic signals for their timing as well as the ways in which adipocytes execute these elegant, interlocking pathways. Our findings support the robust circadian timing of mitochondrial oxidative metabolism, energy metabolism, extracellular matrix (ECM) homeostasis, molecular transport, and transcriptional regulation, while phase alignment is selectively shifted by tissue-specific systemic cues. Together, these data demonstrate that adipocyte circadian output is controlled by cell-intrinsic temporal regulatory mechanisms, while tissue-specific systemic cues modulate phase alignment rather than generate rhythmicity *de novo*.

## RESULTS

### The transcriptome of differentiated adipocytes in culture shows widespread circadian regulation

We isolated pre-adipocytes of inguinal white adipose tissue (iWAT) obtained from Per2::Luc C57BL/6J mice. The pre-adipocytes were then differentiated *in vitro*, and their clocks synchronized via serum shock and forskolin treatment (see Methods and Supplemental Methods for further details). Synchronized Per2::Luc adipocytes exhibited robust ∼24h PER2 oscillations, which were confirmed at the mRNA level by qRT-PCR (Fig. 1A, Fig. S1A-D). Sampling for two replicate time series began at 12 hours post-synchronization (HPS) and continued until 72 HPS. RNA-seq analysis yielded >60M paired-end reads per sample, a depth we chose to detect low-abundance transcripts. To our knowledge, this dataset represents the longest, most deeply sequenced, highest resolution circadian transcriptome for differentiated adipocytes in cell culture to date and has allowed us to detect previously undetected rhythms in CCGs. We performed analysis at the transcript rather than the gene level for protein-coding genes (Fig. S2, Methods, Supplemental Methods). In this dataset, core clock genes were highly circadian, and replicates were almost uniformly in phase (Fig. 1B and Fig. S3). Using ECHO (De Los Santos et al., 2020) and MetaCycle (Wu et al., 2016) with a Benjamini-Hochberg (BH) adjusted p-value ≤ 0.005 and 22-30h period, we identified 4,920 unique circadian transcripts in IVDAs corresponding to 4,346 unique CCGs (21% of the coding transcriptome) (Fig. 1C, D, Supplemental File 1). This represents approximately three times as many CCGs as previously reported for BAT and five as many as was reported for eWAT (cf. Zhang et al., 2014), reflecting use of RNA-seq vs. microarrays, transcript vs. gene level analysis, and ECHO and MetaCycle vs. JTK cycle. ECHO period estimates for rhythmic transcripts in IVDAs are fairly uniform across the detected range (Fig. S4A), the majority of circadian transcripts had BH adj p-values ≤ 0.005, indicating accurate statistical modeling of these genes by ECHO (Fig. S4B), and the distribution of oscillation types shows approximately equal proportions of harmonic and damped oscillations (Fig. S4C) (De Los Santos et al., 2020). As an independent check of our analysis pipeline, as well as to estimate how many CCGs were primary targets of the core clock, we interrogated our list of CCGs against the 6,276 unique BMAL1 target genes identified by iWAT BMAL1 ChIP-seq (Hepler et al., 2022; GEO accession # GSE181443) defining BMAL1-associated targets using a −10 kb to +1 kb window relative to the transcription start site (TSS). The percentage of these CCGs identified as BMAL1 targets does not change significantly as we increase stringency, providing additional validation that our list of CCGs are indeed clock-controlled (Table S1).

**Figure 1.**
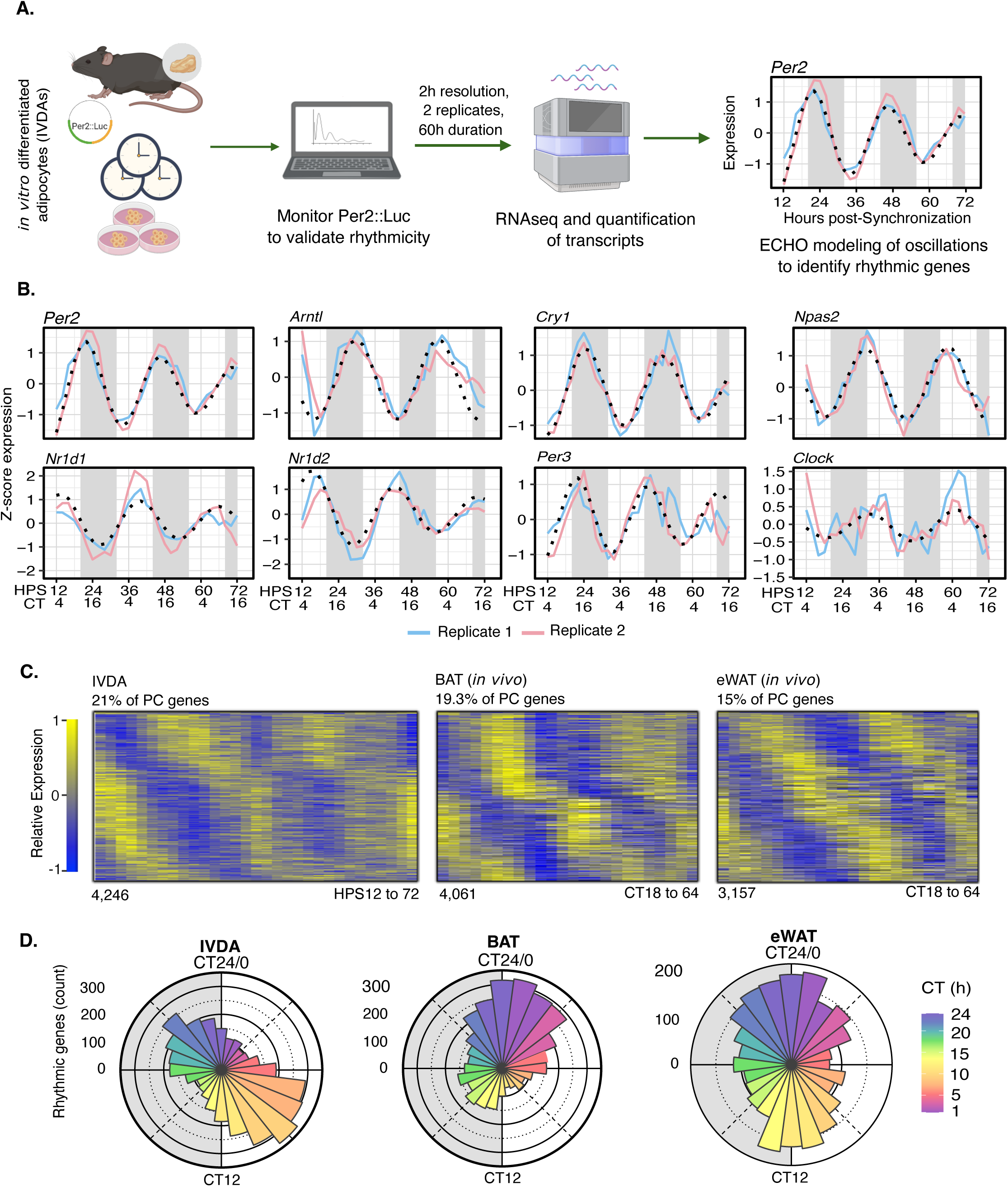
Comparison of the *in vitro* circadian transcriptome with *in vivo* data from BAT and eWAT shows widespread cell-intrinsic circadian regulation. **(A)** Diagram of the analysis of adipocyte circadian regulation. Primary inguinal iWAT adipocytes were harvested from Per2::Luc mice and then differentiated and synchronized *in vitro* via serum shock. Two independent time courses were generated. Replicate one and two were sampled every 2 h over 60 h starting at 12 h post-serum shock. ECHO was used to model oscillations and identify circadianly regulated transcripts. Diagram created in BioRender. Worthen, J. (2026) https://BioRender.com/vf15fbu **(B)** Heatmaps showing the relative expressions of circadian transcripts which oscillated with a period of 22-30h and a BH adj P-value of <0.005 in IVDAs, BAT and eWAT. **(C)** Phase distribution of circadian transcripts in IVDAs and brown adipose tissue (BAT) and epididymal white adipose tissue (eWAT) (*in vivo* microarray data, Zhang et al., 2014) shown in circadian time (CT). **(D)** Oscillation of clock genes in Rep1 (pink) and Rep2 (blue) in IVDAs with the ECHO fitted curve shown in black. X-axis is measured in Z-score and y-axis is shown as hours post-synchronization. IVDA = *In Vitro* Differentiated Adipocytes. PC = Protein-Coding.

### Comparison of the *in vitro* circadian transcriptome with *in vivo* data from BAT and eWAT

A principal goal of this study is to ascertain the relative importance of cell-autonomous versus systemic cues in the rhythmicity of adipocytes. To understand how CCGs identified in our study compare to circadian transcriptomes of mouse adipose tissue influenced by systemic cues, we obtained *in vivo* BAT and eWAT (epididymal white adipose tissue) circadian transcriptome data generated using microarrays (Zhang et al., 2014, GEO accession # GSE54652). We filtered genes to remove transcriptional noise as was done for IVDAs, resulting in a detected protein-coding transcriptome of 11,249 genes for BAT and 11,624 for eWAT (see Methods, Supplemental Files 2 and 3). To ensure accurate comparisons with our dataset, we reanalyzed these datasets using our analysis. ECHO and MetaCycle identified 4,061 rhythmic protein-coding CCGs in BAT and 3,157 in eWAT. This represents 19.3% for BAT and 15% for eWAT, similar to that calculated for IVDAs (Fig. 1C, D) but higher than previously reported (BAT = 8% and eWAT = 4% of protein-coding genes), reflecting the use of ECHO as well as our application of a wider period range compared to Zhang et al. Circadian time enables us to use a common axis to compare phase between organisms with different free-running period lengths. *Per2* expression is robustly circadian in every rhythmic mammalian tissue and consistently peaks about 14h after a dark-to-light transition in adipose tissue from mice (Zhang et al., 2014). To assign an estimate of circadian time (CT) to IVDAs, we used the peak of *Per2* expression (22 HPS) following release from synchronization as a phase reference point, assigning this a phase of CT14 and calculating phases for other genes in reference to *Per2*. By this metric, phases of rhythmic core clock component genes or their paralogs in our IVDAs (e.g., *Npas2* and *Clock*) are within 2 hours of those reported from BAT and eWAT *in vivo* (Zhang et al., 2014) (Table S2, Fig. S5). These results demonstrate that the cell-intrinsic TTFL is functioning similarly in IVDAs when compared to adipose tissue, such that observed differences in the phasing of circadian transcripts are not due to differences in the core oscillator, but rather in the circadian physiology of the IVDAs vs. *in vivo*-sampled adipose.

### Cell-autonomous circadian regulation of a breadth of processes in adipocytes is frequently further sculpted *in vivo* by cell-extrinsic factors

To identify which biological pathways were circadianly regulated and determine whether their timing is dependent on systemic cues, we combined standard enrichment analysis with Phase Set Enrichment Analysis (PSEA), both of which utilized Reactome gene sets (Zhang et al., 2016; Milacic et al., 2024) (Fig. 2). We first conducted a Reactome pathway enrichment analysis of all IVDA CCGs using PANTHER (Thomas et al., 2022; Ashburner et al., 2000; The Gene Ontology Consortium, 2023) and identified RHO, CDC42, RAC, and MIRO GTPase pathways, along with membrane trafficking, transcription, RNA and protein metabolism, cell cycle, and signaling pathways, as significantly enriched for circadian genes (FDR ≤ 0.05) (Fig. 2A). Phase-clustered gene sets are defined as gene sets whose expression peaks occur close together in time. Though PANTHER identifies pathways enriched for circadian genes, it does so without regard to peak phase. To determine whether circadian genes in IVDAs, BAT, and eWAT were phase-clustered, we applied Phase Set Enrichment Analysis (PSEA; Zhang et al., 2016). PSEA Kuiper scores quantify the degree to which circadian gene CT values deviate from a uniform distribution, and pathways with Kuiper q ≤ 0.1 were considered significantly phase-clustered (Methods). Phase clustering strength was further assessed using the vector-average magnitude metric, which reflects temporal cohesiveness within each gene set.

**Figure 2.**
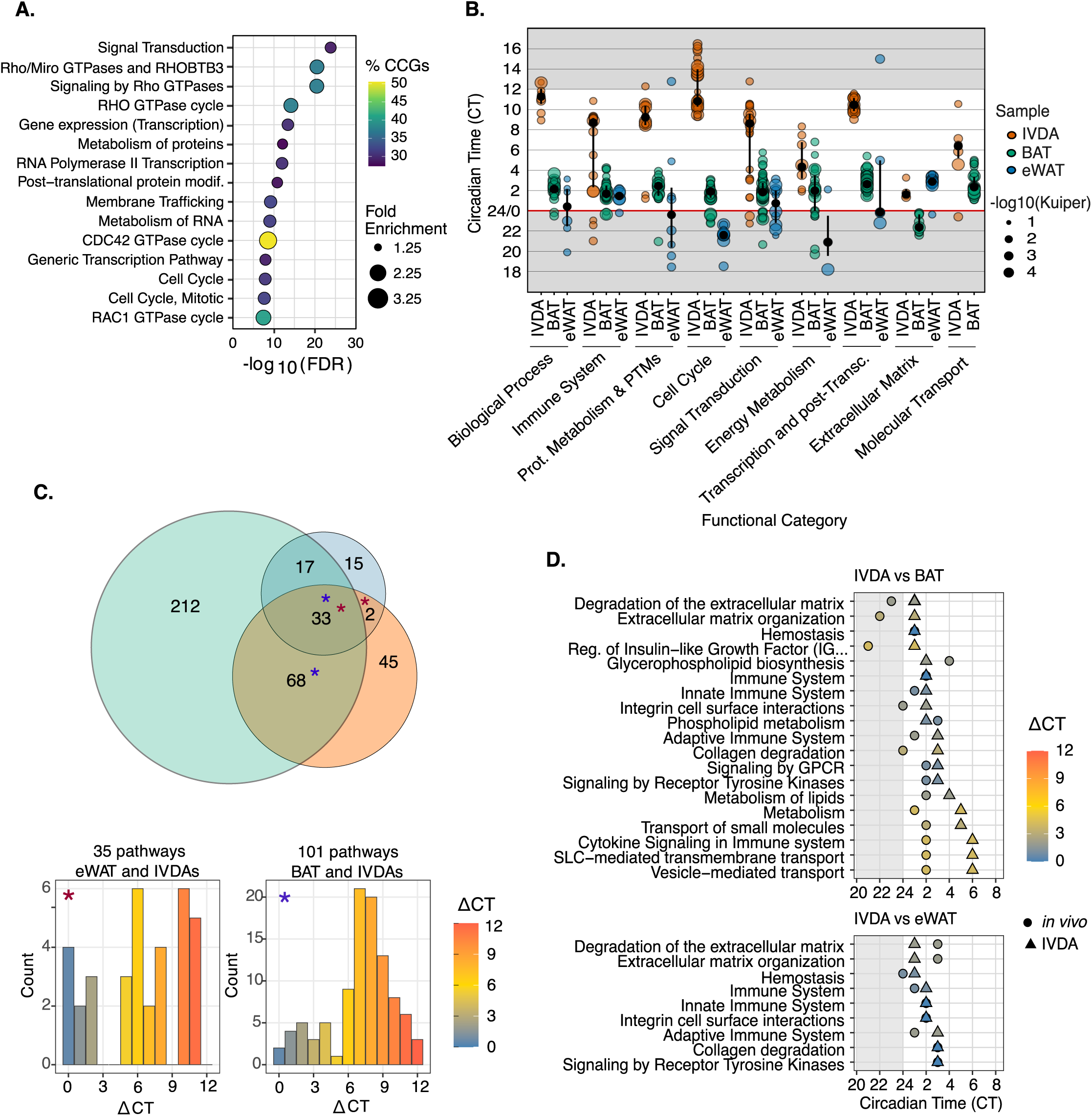
Functional analysis reveals processes that are cell-autonomously circadian yet temporally coordinated by systemic cues. **(A**) Top 15 Reactome top-level groups most significantly enriched for circadian genes by PANTHER (FDR < 0.05) and significantly phase-clustered by PSEA in IVDA. **(B)** Circadian time (CT) of phase-clustered pathways grouped by Reactome top-level process (colored by sample). Point size = -log₁₀ (q value). Gray shading delineates the four regulatory groups described in text. **(C)** Overlap of rhythmic pathways among IVDA, BAT, and eWAT identified by PSEA. Numbers indicate shared pathway counts; histograms show distributions of phase differences (ΔCT) between IVDA and each *in vivo* depot. The IVDA-BAT histogram shows an average 8-9 h phase delay, indicating substantial temporal re-phasing *in vivo*. **(D)** Representative cell-autonomous rhythmic pathways for IVDA vs BAT (left) and eWAT (right) illustrating phase alignment (<4h) within key top-level processes. ECM pathways, although not enriched for circadian genes, were highly phase-clustered in PSEA, suggesting rhythmic coordination of a smaller, coherent gene subset.

For easier interpretation and comparison between *in vitro* and *in vivo* datasets we grouped phase-clustered pathways into functional categories using top-level Reactome terms and visualizing the distribution of peak CTs across tissues (Fig. 2B). Although many pathways are shared between IVDAs, BAT, and eWAT, their circadian timing differs systematically by tissue. Most pathways, including those relating to biological process, immune system, protein metabolism and post-translational modifications (PTMs), cell cycle, signal transduction, and transcription generally peak later in IVDAs than *in vivo* depots, indicating widespread temporal repositioning of circadian programs in IVDAs compared to those *in vivo* (Fig. 2). In contrast, the timing of the energy metabolism pathways was more closely aligned between IVDAs and BAT, while eWAT was selectively shifted. ECM homeostasis pathways exhibit conserved timing across all three tissues, clustering near the day-night transition. The timing of molecular transport pathways was coordinated in IVDAs and BAT with no detected phase-clustering in eWAT. Together, these patterns show that circadian pathway timing depends on both functional category and tissue context, with some programs maintaining similar phase relationships.

To ensure phase clustering isn’t driven by the repeated detection of the same circadian genes across related Reactome terms, we performed a PSEA leading-edge overlap analysis. Pairwise Jaccard similarity among the top 50 most overlapping pathways reveals high gene sharing only among cell-cycle-related pathways; most other functional categories show low overlap (Fig. S6). Phase clustering across diverse biological processes arises from independent circadian gene modules rather than a single dominant driver set, consistent with the high overlap between pathways strongly phase clustered and those highly enriched for circadian genes by PANTHER (Fig. S7). Comparison of vector-average magnitude shows no difference among phase-clustered pathways across all tissues (Fig. S8), indicating that systemic cues primarily modulate pathway timing rather than altering overall temporal cohesiveness.

To measure the degree of cell-autonomous versus systemic circadian regulation of phase-clustered pathways, we compared those detected in IVDAs, BAT, and eWAT and found a high overlap (101 pathways in both BAT and IVDAs, 35 in both eWAT and IVDAs, and 33 shared across all three datasets) (Fig. 2C, top and Fig. S10). A higher resolution representation of all phase-clustered pathways further highlights conservation of circadian pathway architecture with depot-specific phase shifts driven by systemic cues (Fig. S11). BAT-IVDA pathways exhibit a consistent average phase delay of about 8-9 hours, whereas no dominant phase-shift pattern was observed for eWAT (Fig. 2C, bottom and Fig. S11). Pathways conserved and in phase across all three depots were dominated by ECM homeostasis, immune signaling, and receptor tyrosine kinase (RTK) pathways (Fig. 2D and Fig. S10). In contrast, 17 pathways were rhythmic in both BAT and eWAT but not in IVDAs, indicating circadian regulation that depends on systemic inputs rather than adipocyte-intrinsic oscillators (Fig. S12). These pathways were enriched for ECM organization, immune and growth factor signaling, and cholesterol biosynthesis, and peaked during late night to early morning (CT20-4) in both depots. Taken together, these results demonstrate that circadian regulation of adipocyte biology is neither purely intrinsic nor purely systemic but rather emerges from the integration of autonomous oscillators with depot-specific physiological cues that sculpt pathway timing *in vivo*.

### Clock-organized biological work includes energy metabolism

Metabolic processes follow a daily rhythm such that anabolic processes (e.g., glycolysis and lipogenesis) primarily occur during the feeding phase and catabolic processes including lipolysis and fatty acid oxidation (FAO) are engaged during fasting (Liu et al., 2025; Lekkas and Paschos, 2019). Reactome metabolic pathways are strongly enriched for CCGs, with pronounced temporal organization in IVDAs (Fig. 3A, B). Lipid-associated pathways are phase-aligned between BAT and IVDAs but diverge in eWAT, indicating depot-specific regulation of adipocyte metabolism (Fig. 3B and Figs. S9-11).

**Figure 3.**
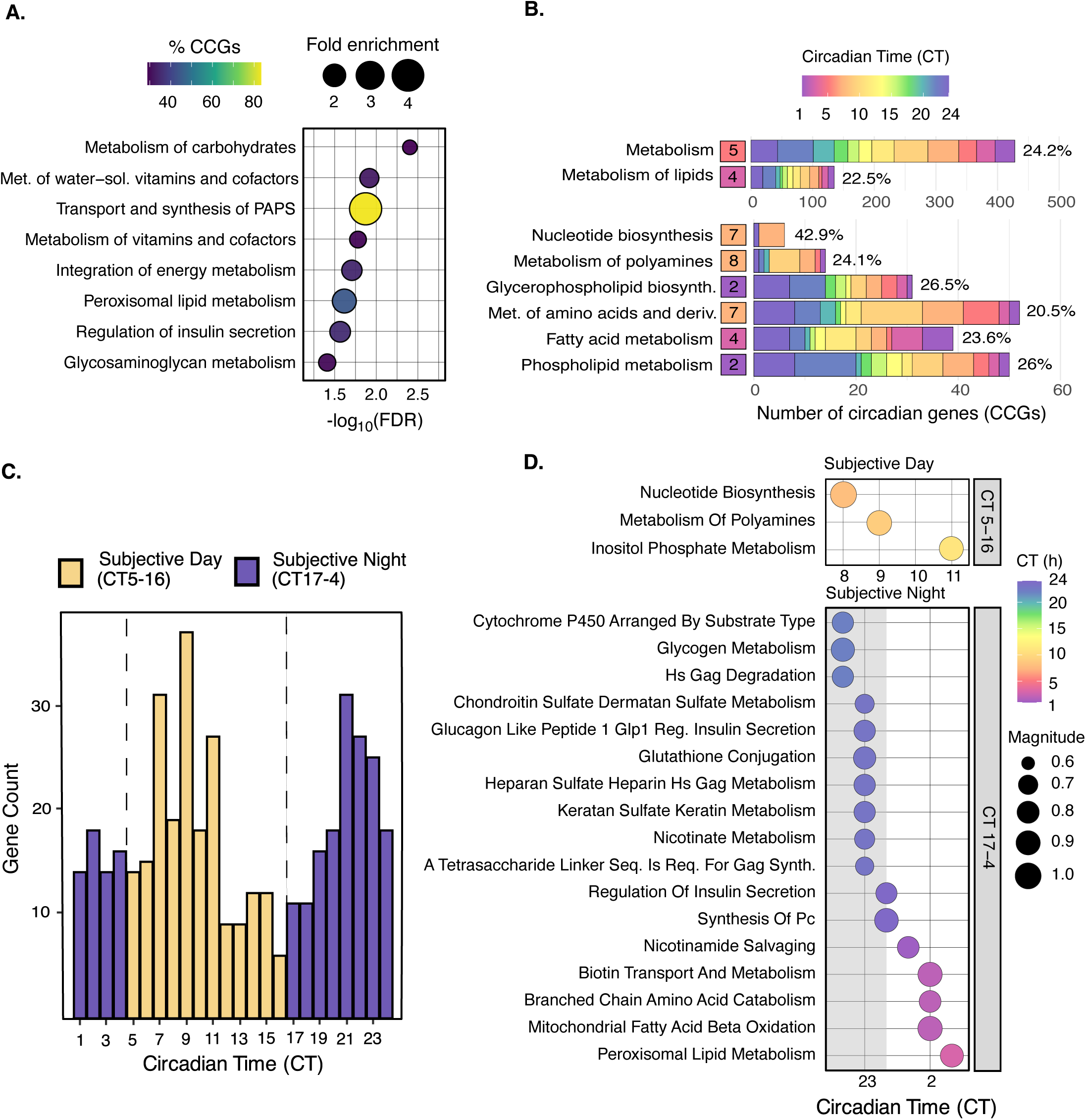
Metabolic sub-pathways display two major circadian waves corresponding to daytime anabolism and nighttime restoration. **(A)** Reactome energy metabolism pathways ranked by circadian enrichment relative to background. The x-axis indicates -log₁₀(FDR) and dot color represents the percentage of circadian genes in each pathhway. Dot size corresponds to fold enrichment. **(B)**. PSEA of Metabolism. Pathways are grouped as Large (> 100 CCGs) or Other (≤ 100 CCGs) and ranked by magnitude. Colored tiles denote mean pathway phase (CT). **(C) His**togram of all Metabolism circadian transcripts showing two phase peaks (CT 5-16 and CT 17-4). (D) PSEA of Metabolism sub-pathways run separately for Peak 1 and Peak 2 genes. Dot plots show the pathways that were Exclusive to either peak 1 (top) or peak 2 (bottom). Color scale = CT; size = magnitude.

The top-level Reactome metabolism pathway shows a significant Kuiper statistic (6x10^-4^) and a very low magnitude (0.14), indicating a bimodal distribution of metabolic CCGs (Fig. 3B, Supplemental File 5). Phase distribution of these genes reveals two dominant peaks: a subjective-day cluster (CT5-16) and a subjective-night cluster (CT17-4) (Fig. 3C). To identify which metabolic processes were specific to each peak, we performed PSEA on the CT5-16 and CT17-4 CCGs independently using only the gene sets which fall under the top-level term of Metabolism. This analysis revealed a clear separation of opposing metabolic processes.

In mice, which are nocturnal, the subjective day overlaps with the resting period (dawn to dusk) whereas the subjective night overlaps with the active period (dusk to dawn). Pathways phase-clustered exclusively within the CT5-16 window are associated with anabolic and biosynthetic processes, consistent with processes happening during the rest-phase. Pathways phase-clustered exclusively within the CT17-4 window are enriched for lipid-centric pathways, including mitochondrial and peroxisomal FAO, phospholipid metabolism, and lipid remodeling which are expected to dominate during the active phase (Hepler et al., 2022) (Fig. 3D). Despite the two windows containing a similar number of CCGs, the nighttime window yielded far more phase-clustered pathways specific to that window suggesting it is the major metabolic window in IVDAs. The CCGs in these pathways were sharply in phase and no genes occur in more than 3 pathways (data not shown), consistent with these being distinct circadianly regulated programs.

Multiple rate-limiting enzymes within these pathways are circadian in IVDAs and in phase with BAT. Enzymes in glycolysis (*Pfkm, Pfkp*) peak late in the circadian cycle (IVDA CT20, 24; BAT CT20, 2). *Acaca*, which catalyzes the first irreversible step in fatty acid synthesis, is circadian and in phase between IVDA and BAT peaking at CT23 and CT20, respectively. Lipolytic enzymes *Pnpla2 and Lipe* peak at CT2 in IVDAs and CT24 in BAT which coincides with the phase of triglyceride mobilization. Fatty acid oxidation is rhythmically controlled by *Cpt1a, Cpt1b, Acadl, Hadha, Hadhb*, and *Acox1* (Liu et al, 2025) which are tightly phase-clustered in the very early morning (IVDA CT2-4; BAT CT2-6). Oxidative metabolism through the TCA cycle (*Ogdh, Idh2, Mdh1*) and branched-chain amino acid degradation (*Bckdha*) also peaks in this window. Additional rate-limiting steps in phospholipid synthesis (*Gpat3*) and NAD⁺ salvage (*Nampt*) peak in phase with lipid handling and mitochondrial function (Supplemental File 1 and 2). Together, these data suggest circadian timing preferentially constrains pathways that support sustained oxidative metabolism, with rhythmic pathways sharing similar timing persisting in cultured adipocytes without systemic cues, underscoring their cell-autonomous rhythmicity.

Pathways associated with the ECM, including integrin and focal adhesion signaling, ECM-receptor interactions, and heparan sulfate and glycosaminoglycan metabolism, show strong phase clustering and peak during the late active to early rest window across IVDAs, BAT, and eWAT (Fig. 2D and Fig. S11). These pathways are in phase with those relating to lipid handling and insulin-related metabolic processes in IVDAs and BAT (Fig. 3D). Adipocyte ECM composition and ECM-cell signaling directly regulate metabolic functions such as insulin-stimulated glucose uptake, adipogenic capacity, and lipid handling, and ECM reprograms adipocyte metabolic phenotypes in a depot-specific manner (Strieder-Barboza et al., 2020; Baker et al., 2017). While circadian regulation of ECM-related pathways has been described in other tissues as a dynamic interface between clocks and cellular function (Dudek et al., 2023), its cell-autonomous circadian regulation has not previously been reported in adipocytes.

### Molecular transport is cell-autonomously circadianly regulated in IVDAs and BAT

Molecular transport pathways were enriched for circadian genes in IVDAs (FDR ≤ 0.05). These include vesicle-mediated transport, membrane trafficking, clathrin-mediated endocytosis, and Golgi-to-ER retrograde transport (Fig. 4A). Intra-Golgi and retrograde trafficking pathways show the strongest enrichment and highest circadian gene content (∼45%), indicating that intracellular transport pathways are strongly circadianly regulated in IVDAs. Consistent with the enrichment results, PSEA analysis also shows phase-clustering of processes related to molecular transport. Large transport pathways (>100 CCGs, Fig. 4B, upper), such as membrane trafficking and vesicle-mediated transport, display more diffuse expression of their CCGs, consistent with continuous transport demand across the circadian day. In contrast, smaller pathways (≤100 CCGs, Fig. 4B, lower) are sharply phase-clustered, peaking primarily during the middle to late subjective day (CT7-14), with plasma lipoprotein assembly and clearance peaking later into the subjective night. Molecular transport pathways are phase-clustered in IVDAs and BAT but not eWAT (Fig. 2B). Although transport pathways peak within a similar circadian window, they are systematically phase-shifted, with BAT peaking near dawn and IVDAs peaking between CT6-11, barring plasma lipoprotein assembly and clearance. Together, these findings indicate that molecular transport follows a cell-intrinsic circadian program whose phase is likely shaped by systemic cues in BAT.

**Figure 4.**
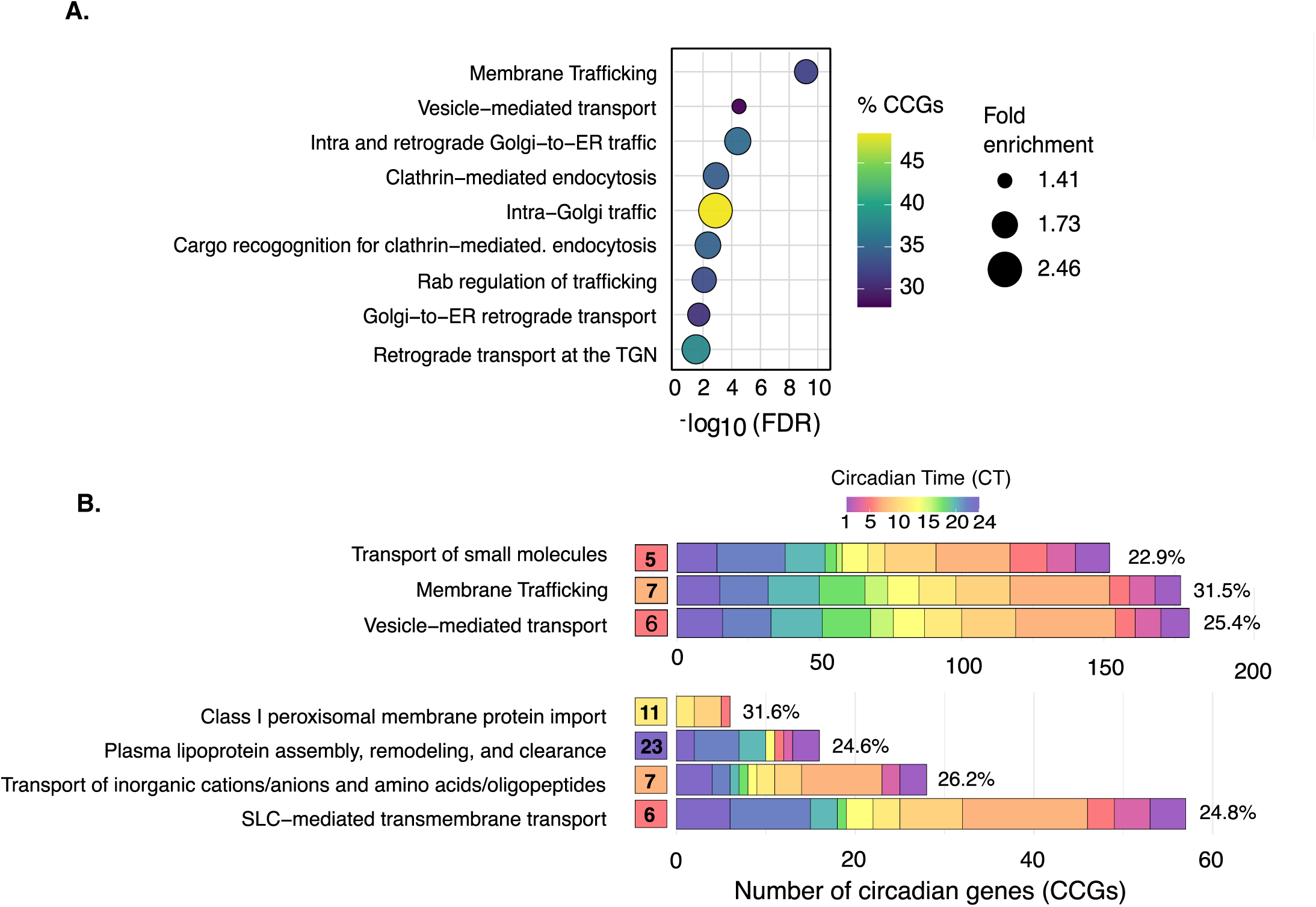
Circadian regulation of molecular transport pathways. **(A)** Enrichment analysis of Reactome transport-related pathways showing fold enrichment versus FDR significance. The x-axis indicates -log₁₀(FDR) and dot color represents the percentage of circadian genes in each pathway. Dot size corresponds to fold enrichment. Analysis done using Panther.org. **(B)** Circadian phase distribution of circadian genes (CCGs) within large (>100 CCGs) and smaller (≤100 CCGs) molecular transport pathways that were determined to be phase clustered (Kuiper q value < 0.1). Bars show the phase distribution of circadian genes color-coded by 2-hour CT bins. Numbers inside tiles indicate the PSEA calculated peak phase for each pathway. Percentages to the right indicate the fraction of total genes in that Reactome pathway that are rhythmic. Large pathways display broad phase coverage, whereas smaller pathways show sharper, phase-specific clustering.

### Transcriptional and post-transcriptional pathways are strongly phase-clustered and peak late-day in IVDAs

Pathway enrichment analysis also revealed that pathways related to transcriptional regulation, chromatin organization, and RNA metabolic processes are strongly enriched for CCGs in IVDAs (Fig. 5A). These pathways span the multiple stages of gene expression, including chromatin-modification, RNA polymerase II-dependent transcription, and rRNA processing and RNA metabolism, including mRNA splicing. Consistent with these enrichment results, PSEA analysis shows phase-clustering of transcription and post-transcriptional pathways (Fig. 5B). These pathways peak sharply within a narrow circadian window around CT10-12, corresponding to the late light (rest) phase, and nearly antiphase to the timing of the same pathways in BAT (Fig. 2B and Figs. S9-11). This phase offset indicates that systemic cues selectively regulate the timing of transcriptional and post-transcriptional processes *in vivo*.

**Figure 5.**
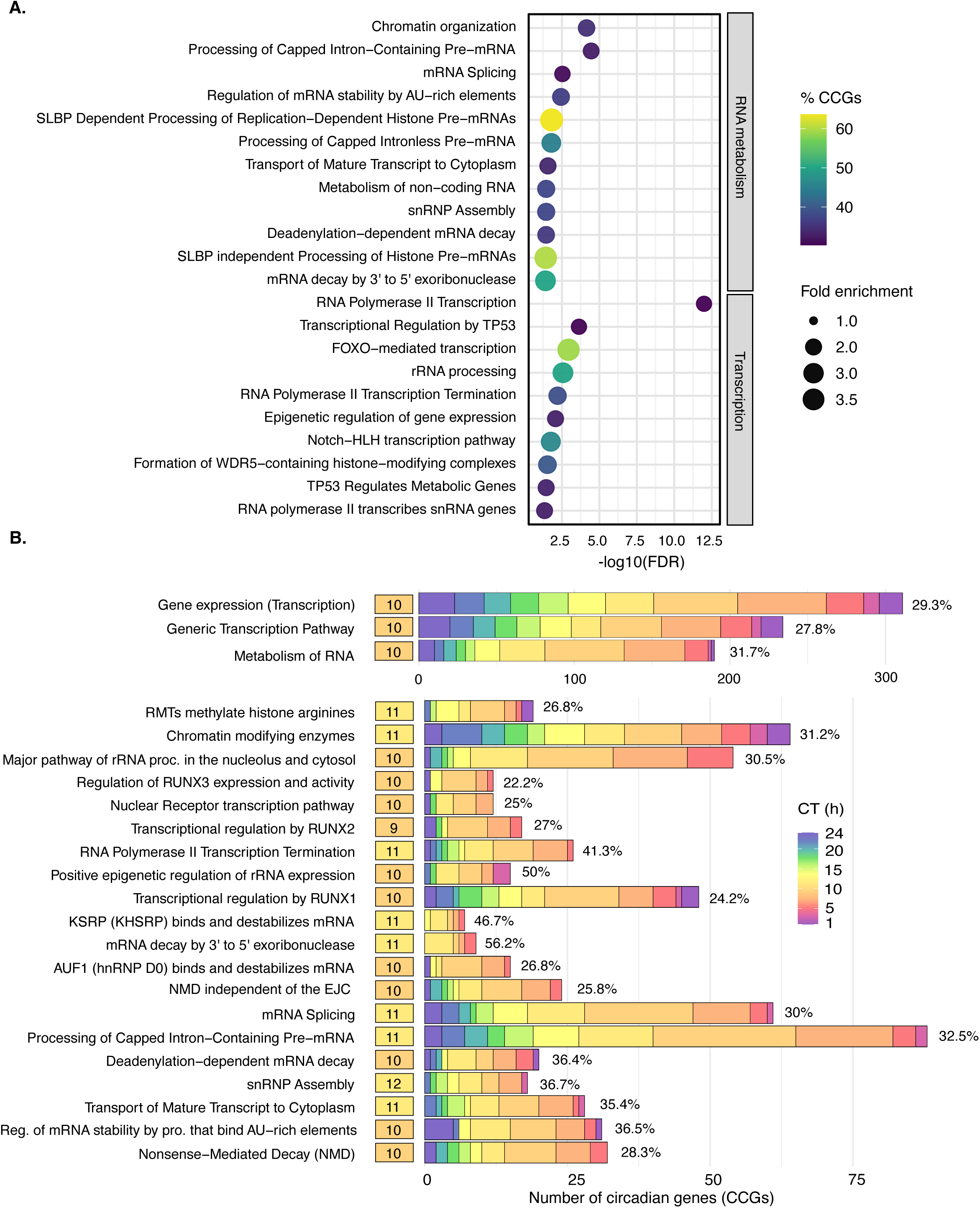
Transcriptional and RNA metabolic pathways are sharply phase-clustered at CT10-12, indicating a coordinated mid-day transcriptional wave in adipocytes. **(A)** Dot plot showing enriched transcriptional and RNA-related pathways. The list of pathways was pruned to reduce redundancy. The x-axis indicates -log₁₀(FDR) and dot color represents the percentage of circadian genes in each pathway. Dot size corresponds to fold enrichment. **(B)** Bar plots showing the number and circadian phase (CT) distribution of circadian genes (CCGs) within each pathway. Pathways are divided into large pathways (>100 CCGs; upper panel) and smaller pathways (≤100 CCGs; lower panel). Bars are colored by CT, and numbers on the right indicate the percentage of total circadian genes represented by each pathway. Pathways are ordered by magnitude.

### Sequential waves of TF-chromatin engagement in rhythmic BAT chromatin

To predict how rhythmicity is regulated at the chromatin level, we analyzed rhythmic BAT ATAC-seq peaks (Hepler et al., 2022) grouped into 4-hour Zeitgeber time (ZT) bins. We then performed motif enrichment analysis (p ≤ 1×10⁻³; BH q ≤ 0.01) on these bins using a curated set of circadian transcription factor (TF) motifs (39 TFs; 42 position weight matrices) (JASPAR, Ovek Baydar et al., 2025) that are phase-aligned between IVDAs and at least one *in vivo* adipose depot (BAT or eWAT) which allowed us to focus on chromatin patterns that reflect circadian regulation that is likely maintained *in vitro*. This analysis revealed distinct temporal clusters of TF-chromatin engagement across the circadian cycle (Fig. 6A). At the global level, our analysis shows that BAT chromatin accessibility oscillates across circadian time, with two prominent peaks in total ATAC-seq signal at ZT9 and ZT17 despite a largely stable number of accessible regions (Fig. S13). These results suggest that it is rhythmic chromatin accessibility, rather than widespread opening or closing of regulatory elements, that provides a chromatin framework for phase-specific TF engagement.

**Figure 6.**
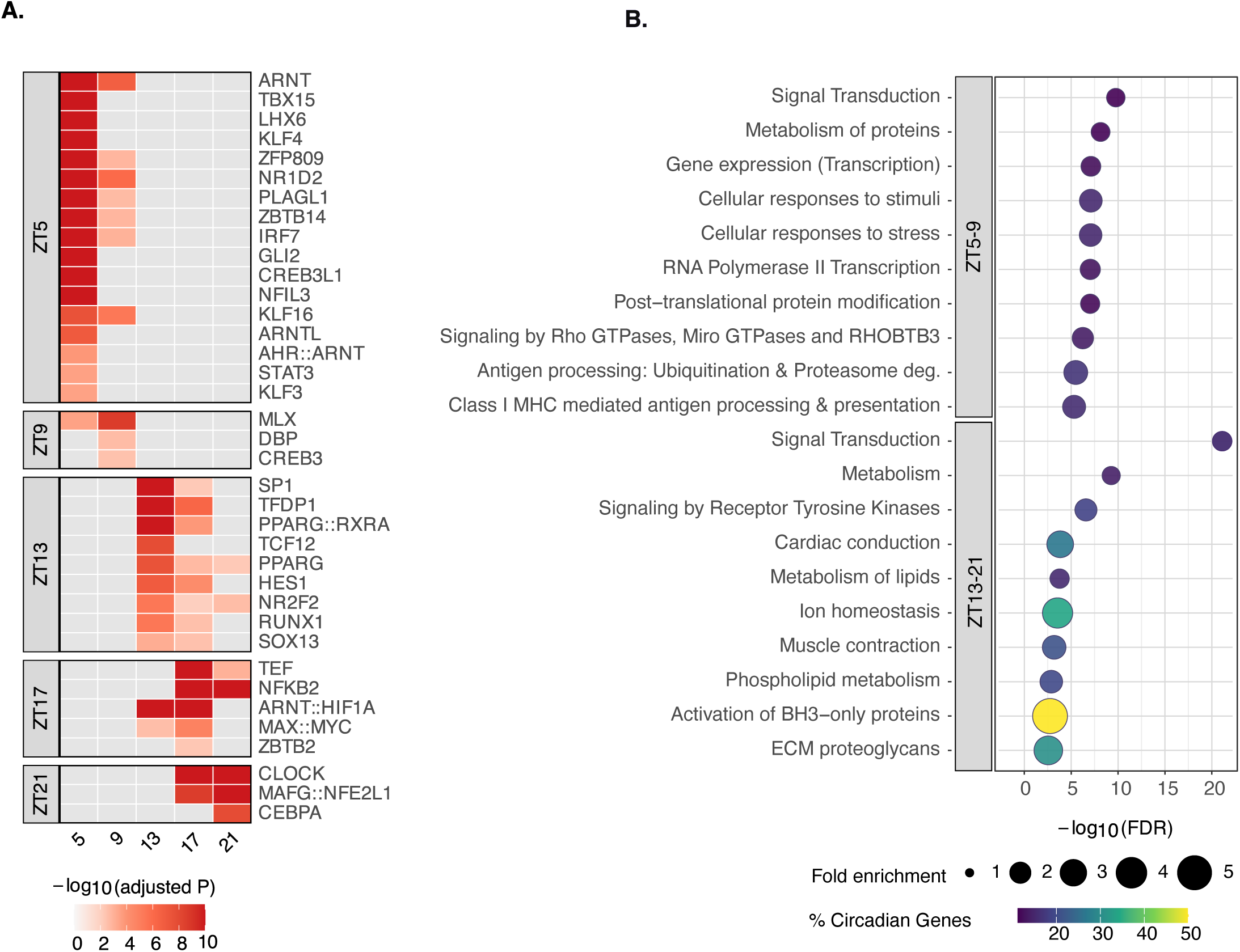
Sequential clusters of circadian transcription factor engagement and pathway enrichment in rhythmic BAT chromatin. **(A)** Heatmap showing significantly enriched circadian transcription factor (TF) motifs (BH-adjusted q < 0.01) across 4-hour Zeitgeber time (ZT) bins in BAT ATAC-seq data. Color indicates -log,(adjusted P-value). Motifs are grouped by the ZT bin at which enrichment is maximal, revealing discrete temporal clusters of TF engagement. **(B)** Reactome pathway enrichment analysis of genes associated with rhythmic BAT chromatin accessibility, summarized across two major temporal clusters (ZT5/9, top and ZT13/17/21, bottom). Dot plots display the top 10 enriched pathways per peak ranked by -log_10_ (FDR). Dot size corresponds to fold enrichment, and color denotes % circadian genes in the pathway.

Motif enrichment in the ZT5-ZT9 bins (early) is dominated by ARNT-family and bHLH-PAS motifs (ARNT, AHR::ARNT, ARNTL), NFIL3/CREB3, and KLF family motifs. The enriched CACGTG E-box motif is annotated as an ARNT-family motif in JASPAR but is also shared by the BMAL1::CLOCK complex. Our analysis shows approximately 70% of the early ZT5 target genes are predicted to be bound by BMAL1 (BMAL1 ChIP-seq peaks in iWAT, Hepler et al., 2022). It is reasonable to assume that this early ARNT enrichment likely reflects clock-dependent chromatin engagement rather than AHR-specific binding, consistent with clock-associated promoter priming near dawn (Menet et al., 2014).

Motifs enriched during the mid-day bin (ZT13) are dominated by nuclear receptor and transcriptional co-regulator motifs, including PPARG::RXRA, NR2F2, TFDP1, and SP1. This enrichment occurs temporally just before the late-day, phase clustered lipid-centric metabolic program identified within the CT17-4 window (Fig. 3D). Motifs enriched during the late-day bins (ZT17-ZT21) are dominated by motifs associated with stress adaptation and feedback regulation, including NFKB2, CLOCK, MAFG::NFE2L1, and CEBPA. This enrichment coincides with the late-day metabolic program identified within the CT17-4 peak. This enrichment profile differs from that reported by Hepler et al. because our analysis was restricted to those TFs that are circadian and in phase between IVDAs and at least one *in vivo* adipose dataset.

Consistent with this temporal organization, enrichment of motifs associated with transcription factors involved in metabolic regulation aligns with the peak expression of several rate-limiting enzymes and key metabolic control points within pathways phase-clustered in the CT17–4 metabolic peak (Fig. 3D). These include enzymes regulating lipid handling, mitochondrial metabolism, energy metabolism, and redox balance: *Lipe, Hadhb*, and *Acox1* (ZT5); *Idh2* and *Nampt* (ZT13); *Pfkp* (ZT17); and *Acadl* (ZT21). These enzymes function within discrete metabolic pathways rather than acting redundantly across multiple pathways, indicating focused circadian regulation at discrete metabolic bottlenecks and coordinated regulation across entire metabolic networks.

To connect transcription factor (TF)-chromatin engagement with biological function, we performed Reactome pathway enrichment analysis of genes associated with rhythmic BAT chromatin accessibility across the two predominant accessibility peaks (ZT5/9 and ZT13-21) (Fig. S14). Early peaks are dominated by pathways related to transcriptional regulation, RNA metabolism, and signaling. In contrast, the late peaks are dominated by pathways relating to lipid metabolism, phospholipid biosynthesis, mitochondrial function, ECM organization, and nuclear receptor signaling (Fig. 6B).

Spatial analysis of the Distance-to-TSS of enriched motifs across the circadian day revealed promoter-proximal motifs dominate early (ZT5-ZT9), whereas distal enhancer-associated motifs and higher motif-to-gene ratios increase later (ZT13-ZT17), before finally declining at the end of the circadian day (ZT21) (Fig. S15). These results begin to construct a model where a cell-autonomous relay of TF-chromatin interactions coordinates the temporal progression from early transcriptional and regulatory programs to later metabolic and structural pathway engagement across the circadian cycle.

## DISCUSSION

To fully understand the role(s) of adipose tissue in mammalian circadian physiology, it is necessary to pursue questions in whole animals, where the full breadth of these systemic actions and interactions can be seen (e.g. Hepler and Bass, 2023; Ribas-Latre et al., 2021; Zhang et al., 2014; Pendergrast et al., 2023). Such studies by design integrate systemic cues such as light/dark, feeding patterns, hormonal regulation, and nutritional status, which influence tissue-specific molecular clocks. However, studying adipose physiology in the context of a whole animal makes it hard to distinguish cell-autonomous regulation from systemic cues, including feedback from one to the other, and thereby potentially difficult to interpret the origins of systemic metabolic mis-regulation. To begin to address this, we generated, to our knowledge, the longest, deepest adipocyte circadian time-course reported to date. This, in combination with ECHO and MetaCycle, allowed us to identify more transcripts and clock-regulated transcripts than previously possible in adipocytes, as well as more rhythmic genes in BAT and eWAT. Using these rhythmic genes, PSEA analyses allowed insights into the nature of cell-autonomous circadian regulation of adipocyte transcriptional output. Beyond this, while it is plain to see that cell-autonomous rhythmicity in IVDAs and *in vivo* is quite pervasive, an attempt to dissect all of this output for all processes risked saying very little about very much. To avoid this and to gain insight into regulatory processes with the goal of revealing cell-autonomous vs. systemic regulation, we concentrated on pathways governing transcriptional regulation, RNA metabolism, molecular transport, ECM, and energy metabolism.

We identified far more temporally regulated processes in BAT vs IVDAs or eWAT. This increase may suggest that the clock is involved in the regulation of more processes in metabolically active tissues so as to offer a higher degree of metabolic flexibility. There is also a daily rhythm in feeding and body temperature to drive metabolic processes *in vivo*. An organism may benefit from increased circadian regulation in metabolically active tissues to offer a higher degree of metabolic flexibility due in part to fluctuations in the availability of nutrients.

Agreeing with this observation, many phase-clustered pathways detected by PSEA were shared between IVDAs and BAT, whereas overlap with eWAT was less abundant, which suggests depot-specific differences in the timing of their regulation (Fig. 2C). The circadian clock empowers varying degrees of metabolic flexibility across tissues (Thurley et al., 2017). This is the other side of the same molecular coin; a highly circadianly regulated tissue, like BAT compared to eWAT may be more susceptible to circadian disruption, given that misalignment by systemic cues is proportional to its metabolic flexibility.

Together, the BAT chromatin accessibility data support the transcriptional similarities observed between IVDAs and BAT and provide a regulatory framework for these shared rhythms. Rhythmic BAT chromatin accessibility follows temporally ordered patterns that align with phase-clustered transcriptional and metabolic programs also present in IVDAs. The correspondence between TF-chromatin engagement, pathway enrichment, and the timing of rate-limiting metabolic enzymes suggests that core aspects of oxidative metabolism, lipid handling, and mitochondrial function are governed by cell-autonomous chromatin programs that are preserved in BAT *in vivo*. These findings indicate that IVDAs capture key regulatory features of BAT metabolism, supporting the idea that shared intrinsic circadian mechanisms underlie their similar transcriptional and metabolic organization, while systemic cues primarily modulate timing rather than defining these programs. Differences between our motif enrichment analysis and prior results reported by Hepler et al. reflect the use of different motif enrichment algorithms and our input list of motifs being restricted to those for TFs that are circadian and in phase between IVDAs and at least one in the *in vivo* dataset. Despite these differences, both analyses converge on a temporally ordered chromatin landscape linking TF engagement with metabolic and mitochondrial pathways in BAT.

## MATERIALS AND METHODS

### Animals and cell culture

The inguinal stromal vascular fraction (SVF) was isolated from 8-to 12-week-old homozygous Per2::Luc C57BL/6J mice (N=4) housed at 21±2°C under a 12:12h light/dark cycle and free access to normal chow (PicoLab Verified - 75 IF; 23.3% PRO, 12.9% FAT, 63.7% CHO). Isolated preadipocytes were cultured in "differentiation media" before transfer to "maintenance media". Cells were synchronized for 2-2.5 hours with 50% "base media", 50% FBS (Gibco, lot# 2102306), and 10µM forskolin, then switched to "maintenance media" without dexamethasone and containing 0.01% d-luciferin (100 nM, GoldBio #LUCK-100). Rhythms in luciferase activity were monitored with a Lumicycle to confirm clocks were synchronous across samples. Quantitative PCR of *Per2* was used to confirm synchronicity. See Supplemental Methods and Fig. S1 for further details. Sampling for two replicate time series began at 12 hours post-synchronization (HPS) to avoid artifacts immediately following serum shock (Balsalobre et al., 1998) and continued every 2h with the final sample collected at 72 HPS.

### RNA extraction, RNA-sequencing and quantification

Total RNA was extracted from cells using the Qiagen RNeasy Lipid Tissue Mini Kit (Cat #74804, Qiagen) according to manufacturer’s protocol. Directional RNA-seq libraries were prepared with ≥ 2µg RNA using rRNA depletion and sequenced to a depth of at least 60M PE reads/sample to allow transcript isoform-level analyses, following the guidelines for detecting low-abundance and isoform-specific circadian transcripts (Li et al., 2015). Reads mapping to any residual ribosomal rRNA were removed using SortMeRNA (Kopylova et al., 2012) and remaining reads were mapped to the *Mus musculus* GRCm38 reference genome and quantified using RSEM version 1.3.1 to produce expression in transcripts per million (TPM). See Supplemental Methods for further details.

### Filtering of transcripts

To reduce noise and exclude unexpressed and non-productive transcripts from our analysis, we filtered transcripts to an average Count Per Million (CPM avg) ≥ 10 across all values from replicates and limited our analysis to transcripts that map to protein-coding genes (Fig. S2). Noncoding transcripts were not analyzed and will be the focus of a future manuscript. This filtering process yielded 29,020 transcripts mapping to 13,601 unique genes. For accurate comparison, we similarly filtered BAT and epididymal WAT (eWAT) microarray data from Zhang et al., 2014 to remove transcriptional noise and included only expressed genes. This resulted in a detected protein-coding transcriptome of 11,249 genes for BAT and 11,624 for eWAT. For both IVDAs and *in vivo* datasets, genes under these cutoffs were considered unexpressed and excluded from analysis.

### Preprocessing and rhythmicity calling

The data for IVDAs were preprocessed with LIMBR (Crowell et al., 2019) to remove batch effects. After filtering out noise and unexpressed transcripts (see Fig. S2 and Supplemental Methods), screening for rhythmicity was performed using Extended Circadian Harmonic Oscillator (ECHO; De los Santos et al., 2020). Candidate circadian genes were classified as rhythmic using ECHO based on the following criteria: one or more of their transcripts oscillating with a period of 22-30h; oscillation type of harmonic, damped or forced; Benjamini-Hochberg -adjusted p-value ≤ 0.005. This stringent cutoff ensures only the most strongly circadian transcripts were included in our analysis. If more than one transcript passed the requirements for CCG calling the transcript with the lowest BH-adj p-value was selected as the gene’s representative transcript. We also used MetaCycle’s “meta2d” (Wu et al., 2016) as a complementary algorithm for rhythmicity detection, using the same rhythmicity cutoffs as described above.

### Use of PANTHER and PSEA

Methods and parameters applied for use of PANTHER (gene set enrichment analysis) to identify pathways positively enriched for circadian genes and PSEA (to detect phase-clustered pathways) are fully described in Methods and Supplemental File 4 and 5.

## Acknowledgements

We would like to thank Shannon Soucy, Tim Sullivan and Owen Wilkins at the Dartmouth Analytics Core for their consultation on data analysis and interpretation. We would also like to thank Joseph Takahashi who gifted the Per2::Luc mouse prior to the SNP-assisted Speed Congenic crossing at Dartmouth. We would also like to thank Bruce Spiegelman at Harvard Medical School for his valuable advice and guidance in this project.

## Author Contributions

Conceived and designed the experiments: JMW, AMF, JJL, JCD. Performed the experiments: JMW and AMF. Analyzed the data: JMW, AMF, PAD. Wrote the paper: JMW, AMF, PAD and JCD.

## Data availability

All raw RNA-sequencing data generated in this study have been submitted to the NCBI Gene Expression Omnibus (GEO) repository under accession number GSE202686. eWAT (epididymal white adipose tissue) and BAT (brown adipose tissue) circadian transcriptome data generated by Zhang et al., 2014 using microarrays can be found under GEO accession GSE54652. BMAL1 ChIP-seq data generated by Hepler et al., 2022 can be found in the GEO repository (GSE181443).

## Supplemental Files

Supplemental_Methods.pdf

Supplemental_Figures.pdf

Supplemental_File_S1.xlsx

Supplemental_File_S2.xlsx

Supplemental_File_S3.xlsx

Supplemental_File_S4.xlsx

Supplemental_File_S5.xlsx

Supplemental_Table_S1.docx

Supplemental_Table_S2.docx

## CITATIONS

Acosta-Rodríguez V, Rijo-Ferreira F, Izumo M, Xu P, Wight-Carter M, Green CB, Takahashi JS. 2022. Circadian alignment of early onset caloric restriction promotes longevity in male C57BL/6J mice. Science Jun 10;376(6598):1192–1202. doi: 10.1126/science.abk0297. Epub 2022 May 5. PMID: 35511946; PMCID: PMC9262309.

Acosta-Rodríguez V, Rijo-Ferreira F, van Rosmalen L, Izumo M, Park N, Joseph C, Hepler C, Thorne AK, Stubblefield J, Bass J, Green CB, Takahashi JS. 2024. Misaligned feeding uncouples daily rhythms within brown adipose tissue and between peripheral clocks. Cell Rep. Aug 27;43(8):114523. doi: 10.1016/j.celrep.2024.114523. Epub 2024 Jul 23. PMID: 39046875; PMCID: PMC12223410.

Allada R, Bass J. Circadian Mechanisms in Medicine. 2021. N Engl J Med Feb 11;384(6):550–561. doi: 10.1056/NEJMra1802337. PMID: 33567194; PMCID: PMC8108270.

Ashburner M, Ball CA, Blake JA, Botstein D, Butler H, Cherry JM, Davis AP, Dolinski K, Dwight SS, Eppig JT, et al. 2000. Gene ontology: tool for the unification of biology. The Gene Ontology Consortium. Nat Genet May;25(1):25–9. doi: 10.1038/75556. PMID: 10802651; PMCID: PMC3037419.

Baker NA, Muir LA, Washabaugh AR, Neeley CK, Chen SY, Flesher CG, Vorwald J, Finks JF, Ghaferi AA, Mulholland MW, et al. 2017. Diabetes-Specific Regulation of Adipocyte Metabolism by the Adipose Tissue Extracellular Matrix. J Clin Endocrinol Metab Mar 1;102(3):1032–1043. doi: 10.1210/jc.2016-2915. PMID: 28359093; PMCID: PMC5460687.

Balsalobre A, Damiola F, Schibler U. 1998. A serum shock induces circadian gene expression in mammalian tissue culture cells. Cell Jun 12;93(6):929–37. doi: 10.1016/s0092-8674(00)81199-x. PMID: 9635423.

Bass J. 2012. Circadian topology of metabolism. Nature Nov 15;491(7424):348–56. doi:10.1038/nature11704. PMID: 23151577.

Bertholet AM, Kazak L, Chouchani ET, Bogaczyńska MG, Paranjpe I, Wainwright GL, Bétourné A, Kajimura S, Spiegelman BM, Kirichok Y. 2017. Mitochondrial Patch Clamp of Beige Adipocytes Reveals UCP1-Positive and UCP1-Negative Cells Both Exhibiting Futile Creatine Cycling. Cell Metab Apr 4;25(4):811–822.e4. doi: 10.1016/j.cmet.2017.03.002. PMID: 28380374; PMCID: PMC5448977.

Bunk J, Hussain MF, Delgado-Martin M, Samborska B, Ersin M, Shaw A, Rahbani JF, Kazak L. 2025. The Futile Creatine Cycle powers UCP1-independent thermogenesis in classical BAT. Nat Commun Apr 4;16(1):3221. doi: 10.1038/s41467-025-58294-4. PMID: 40185737; PMCID: PMC11971250.

Cannon B, Nedergaard J. 2004. Brown adipose tissue: function and physiological significance. Physiol Rev Jan;84(1):277–359. doi: 10.1152/physrev.00015.2003. PMID: 14715917.

Cao X, Yang Y, Selby CP, Liu Z, Sancar A. 2021. Molecular mechanism of the repressive phase of the mammalian circadian clock. Proc Natl Acad Sci U S A Jan 12;118(2):e2021174118. doi:10.1073/pnas.2021174118. Epub 2020 Dec 21. PMID: 33443219; PMCID: PMC7812753.

Corvera S, Rajan A, Townsend KL, Shamsi F, Wu J, Svensson KJ, Zeltser LM, Collins S, Reis T, Tseng YH, et al. 2026. Advances in Adipose Tissue Biology. Endocr Rev Jan 13;47(1):75–92. doi: 10.1210/endrev/bnaf032. PMID: 41071598.

Crowell AM, Greene CS, Loros JJ, Dunlap JC. 2019. Learning and Imputation for Mass-spec Bias Reduction (LIMBR). Bioinformatics May 1;35(9):1518–1526. doi:10.1093/bioinformatics/bty828. PMID: 30247517; PMCID: PMC6499252.

Cypess AM, Cannon B, Nedergaard J, Kazak L, Chang DC, Krakoff J, Tseng YH, Schéele C, Boucher J, Petrovic N, et al. 2025. Emerging debates and resolutions in brown adipose tissue research. Cell Metab Jan 7;37(1):12-33. doi: 10.1016/j.cmet.2024.11.002. Epub 2024 Dec 6. PMID: 39644896; PMCID: PMC11710994.

De Los Santos H, Collins EJ, Mann C, Sagan AW, Jankowski MS, Bennett KP, Hurley JM. 2020. ECHO: an application for detection and analysis of oscillators identifies metabolic regulation on genome-wide circadian output. Bioinformatics Feb 1;36(3):773–781. doi:10.1093/bioinformatics/btz617. PMID: 31384918; PMCID: PMC7523678.

Deota S, Pendergast JS, Kolthur-Seetharam U, Esser KA, Gachon F, Asher G, Dibner C, Benitah SA, Escobar C, Muoio DM, et al. 2025. The time is now: accounting for time-of-day effects to improve reproducibility and translation of metabolism research. Nat Metab Mar;7(3):454–468. doi: 10.1038/s42255-025-01237-6. Epub 2025 Mar 17. PMID: 40097742; PMCID: PMC12584160.

Dudek M, Swift J, Meng QJ. 2023. The circadian clock and extracellular matrix homeostasis in aging and age-related diseases. Am J Physiol Cell Physiol Jul 1;325(1):C52–C59. doi:10.1152/ajpcell.00122.2023. Epub 2023 May 29. PMID: 37246635; PMCID: PMC10281784.

Fekry B, Eckel-Mahan K. 2022. The circadian clock and cancer: links between circadian disruption and disease Pathology. J Biochem May 11;171(5):477–486. doi: 10.1093/jb/mvac017. PMID:35191986.

Hasan N, Nagata N, Morishige JI, Islam MT, Jing Z, Harada KI, Mieda M, Ono M, Fujiwara H, Daikoku T, et al. 2021. Brown adipocyte-specific knockout of Bmal1 causes mild but significant thermogenesis impairment in mice. Mol Metab Jul;49:101202. doi: 10.1016/j.molmet.2021.101202. Epub 2021 Mar 3. PMID: 33676029; PMCID: PMC8042177.

Hepler C, Bass J. 2023. Circadian mechanisms in adipose tissue bioenergetics and plasticity. Genes Dev Jun 1;37(11-12):454–473. doi: 10.1101/gad.350759.123. Epub 2023 Jun 26. PMID:37364987; PMCID: PMC10393195.

Hepler C, Weidemann BJ, Waldeck NJ, Marcheva B, Cedernaes J, Thorne AK, Kobayashi Y, Nozawa R, Newman MV, Gao P, et al. 2022. Time-restricted feeding mitigates obesity through adipocyte thermogenesis. Science Oct 21;378(6617):276–284. doi: 10.1126/science.abl8007. Epub 2022 Oct 20. PMID: 36264811; PMCID: PMC10150371.

Hurley JM, Loros JJ, Dunlap JC. 2016. The circadian system as an organizer of metabolism. Fungal Genet Biol May;90:39–43. doi: 10.1016/j.fgb.2015.10.002. Epub 2015 Oct 20. PMID:26498192; PMCID: PMC4818683.

Inagaki T, Sakai J, Kajimura S. 2016. Transcriptional and epigenetic control of brown and beige adipose cell fate and function. Nat Rev Mol Cell Biol Aug;17(8):480–95. doi:10.1038/nrm.2016.62. Epub 2016 Jun 2. Erratum in: Nat Rev Mol Cell Biol. 2017Aug;18(8):527. doi: 10.1038/nrm.2017.72. PMID: 27251423; PMCID: PMC4956538.

Kettner NM, Mayo SA, Hua J, Lee C, Moore DD, Fu L. 2015. Circadian Dysfunction Induces Leptin Resistance in Mice. Cell Metab Sep 1;22(3):448–59. doi: 10.1016/j.cmet.2015.06.005. Epub 2015 Jul 9. PMID: 26166747; PMCID: PMC4558341.

Kiehn JT, Tsang AH, Heyde I, Leinweber B, Kolbe I, Leliavski A, Oster H. 2017. Circadian Rhythms in Adipose Tissue Physiology. Compr Physiol Mar 16;7(2):383–427. doi:10.1002/cphy.c160017. PMID: 28333377.

Kopylova E, Noé L, Touzet H. 2012. SortMeRNA: fast and accurate filtering of ribosomal RNAs in metatranscriptomic data. Bioinformatics Dec 15;28(24):3211–3217. doi:10.1093/bioinformatics/bts611. PMID: 23071270.

Lekkas D, Paschos GK. 2019. The circadian clock control of adipose tissue physiology and metabolism. Auton Neurosci Jul;219:66–70. doi: 10.1016/j.autneu.2019.05.001. Epub 2019 May 8. PMID: 31122604.

Li J, Grant GR, Hogenesch JB, Hughes ME. 2015. Considerations for RNA-seq analysis of circadian rhythms. Methods Enzymol 2015;551:349–67. doi: 10.1016/bs.mie.2014.10.020. Epub 2014 Dec26. PMID: 25662464.

Liu H, Wang S, Wang J, Guo X, Song Y, Fu K, Gao Z, Liu D, He W, Yang LL. 2025. Energy metabolism in health and diseases. Signal Transduct Target Ther Feb 18;10(1):69. doi: 10.1038/s41392-025-02141-x. PMID: 39966374; PMCID: PMC11836267.

Loft A, Emont MP, Weinstock A, Divoux A, Ghosh A, Wagner A, Hertzel AV, Maniyadath B, Deplancke B, Liu B, et al. 2025. Towards a consensus atlas of human and mouse adipose tissue at single-cell resolution. Nat Metab May;7(5):875–894. doi: 10.1038/s42255-025-01296-9. Epub 2025 May 13. PMID: 40360756; PMCID: PMC12707904.

Malik DM, Rhoades SD, Zhang SL, Sengupta A, Barber A, Haynes P, Arnadottir ES, Pack A, Kibbey RG, Kain P, et al. 2024. Glucose Challenge Uncovers Temporal Fungibility of Metabolic Homeostasis over a day:night cycle. bioRxiv [Preprint]. Aug 21:2023.10.30.564837. doi: 10.1101/2023.10.30.564837. PMID: 37961230; PMCID: PMC10634956.

Maniyadath B, Zhang Q, Gupta RK, Mandrup S. 2023. Adipose tissue at single-cell resolution. Cell Metab 35(3):386–413. Mar 7. doi: 10.1016/j.cmet.2023.02.002. PMID: 36870444; PMCID: PMC10021468.

Menet JS, Pescatore S, Rosbash M. 2014. CLOCK:BMAL1 is a pioneer-like transcription factor. Genes Dev Jan 1;28(1):8–13. doi: 10.1101/gad.228536.113. PMID: 24395244; PMCID:PMC3894415.

Milacic M, Beavers D, Conley P, Gong C, Gillespie M, Griss J, Haw R, Jassal B, Matthews L, May B, et al. 2024. The Reactome Pathway Knowledgebase 2024. Nucleic Acids Res Jan 5;52(D1):D672–D678. doi: 10.1093/nar/gkad1025. PMID: 37941124; PMCID: PMC10767911.

Narasimamurthy R, Virshup DM. 2021. The phosphorylation switch that regulates ticking of the circadian clock. Mol Cell Mar 18;81(6):1133–1146. doi: 10.1016/j.molcel.2021.01.006. Epub 2021 Feb 4. PMID: 33545069.

Neufeld-Cohen A, Robles MS, Aviram R, Manella G, Adamovich Y, Ladeuix B, Nir D, Rousso-Noori L, Kuperman Y, Golik M, et al. 2016. Circadian control of oscillations in mitochondrial rate-limiting enzymes and nutrient utilization by PERIOD proteins. Proc Natl Acad Sci U S A Mar 22;113(12):E1673-82. doi: 10.1073/pnas.1519650113. Epub 2016 Feb 9. PMID: 26862173; PMCID: PMC4812734.

Ovek Baydar D, Rauluseviciute I, Aronsen DR, Blanc-Mathieu R, Bonthuis I, de Beukelaer H, Ferenc K, Jegou A, Kumar V, Lemma RB, et al. 2026. JASPAR 2026: expansion of transcription factor binding profiles and integration of deep learning models. Nucleic Acids Res Jan 6;54(D1):D184-D193. doi: 10.1093/nar/gkaf1209. PMID: 41325984; PMCID: PMC12807658.

Pendergrast LA, Lundell LS, Ehrlich AM, Ashcroft SP, Schönke M, Basse AL, Krook A, Treebak JT, Dollet L, Zierath JR. 2023. Time of day determines postexercise metabolism in mouse adipose tissue. Proc Natl Acad Sci U S A Feb 21;120(8):e2218510120. doi: 10.1073/pnas.2218510120. Epub 2023 Feb 13. PMID: 36780527; PMCID: PMC9974500.

Philpott JM, Freeberg AM, Park J, Lee K, Ricci CG, Hunt SR, Narasimamurthy R, Segal DH, Robles R, Cai Y, Tripathi S, McCammon JA, Virshup DM, Chiu JC, Lee C, Partch CL. 2023. PERIOD phosphorylation leads to feedback inhibition of CK1 activity to control circadian period. Mol Cell May 18;83(10):1677–1692.e8. doi: 10.1016/j.molcel.2023.04.019. PMID: 37207626; PMCID: PMC11684667.

Puigserver P, Wu Z, Park CW, Graves R, Wright M, Spiegelman BM. 1998. A cold-inducible coactivator of nuclear receptors linked to adaptive thermogenesis. Cell Mar 20;92(6):829–39. doi: 10.1016/s0092-8674(00)81410-5. PMID: 9529258.

Rahbani JF, Bunk J, Lagarde D, Samborska B, Roesler A, Xiao H, Shaw A, Kaiser Z, Braun JL, Geromella MS, et al. 2024. Parallel control of cold-triggered adipocyte thermogenesis by UCP1 and CKB. Cell Metab Mar 5;36(3):526–540.e7. doi: 10.1016/j.cmet.2024.01.001. Epub 2024 Jan 24. PMID: 38272036.

Rasmussen ES, Takahashi JS, Green CB. 2022. Time to target the circadian clock for drug discovery. Trends Biochem Sci Sep;47(9):745–758. doi: 10.1016/j.tibs.2022.04.009. Epub 2022 May 13. PMID: 35577675; PMCID: PMC9378619.

Ribas-Latre A, Santos RB, Fekry B, Tamim YM, Shivshankar S, Mohamed AMT, Baumgartner C, Kwok C, Gebhardt C, Rivera A, et al. 2021. Cellular and physiological circadian mechanisms drive diurnal cell proliferation and expansion of white adipose tissue. Nat Commun Jun 9;12(1):3482. doi: 10.1038/s41467-021-23770-0.

Ricci CG, Philpott JM, Torgrimson MR, Freeberg AM, Narasimamurthy R, Pécora de Barros E, Amaro R, Virshup DM, McCammon JA, Partch CL. 2025. Markovian state models uncover casein kinase 1 dynamics that govern circadian period. Biophys J Nov 18;124(22):4034–4048. doi: 10.1016/j.bpj.2025.09.022. Epub 2025 Sep 18. PMID: 40968534; PMCID: PMC12709424.

Roh HC, Tsai LTY, Shao M, Tenen D, Shen Y, Kumari M, Lyubetskaya A, Jacobs C, Dawes B, Gupta RK, et al. 2018. Warming Induces Significant Reprogramming of Beige, but Not Brown, Adipocyte Cellular Identity. Cell Metab May 1;27(5):1121–1137.e5. doi: 10.1016/j.cmet.2018.03.005. Epub 2018 Apr 12. PMID: 29657031; PMCID: PMC5932137.

Rosen ED, Spiegelman BM. 2006. Adipocytes as regulators of energy balance and glucose homeostasis. Nature Dec 14;444(7121):847–53. doi: 10.1038/nature05483. PMID:17167472; PMCID: PMC3212857.

Sakers A, De Siqueira MK, Seale P, Villanueva CJ. 2022. Adipose-tissue plasticity in health and disease. Cell Feb 3;185(3):419–446. doi: 10.1016/j.cell.2021.12.016. PMID: 35120662;PMCID: PMC11152570.

Samanta S, Ali SA. 2022. Impact of circadian clock dysfunction on human health. Explor neurosci 1(1), Article 1. 10.37349/en.2022.00002.

Scheer FA, Hilton MF, Mantzoros CS, Shea SA. 2009. Adverse metabolic and cardiovascular consequences of circadian misalignment. Proc Natl Acad Sci U S A Mar 17;106(11):4453–8. doi: 10.1073/pnas.0808180106. Epub 2009 Mar 2. PMID: 19255424; PMCID: PMC2657421.

Schrader LA, Ronnekleiv-Kelly SM, Hogenesch JB, Bradfield CA, Malecki KM. 2024. Circadian disruption, clock genes, and metabolic health. J Clin Invest Jul 15;134(14):e170998. doi: 10.1172/JCI170998. PMID: 39007272; PMCID: PMC11245155.

Smyllie NJ, Koch AA, Adamson AD, Patton AP, Johnson A, Bagnall JS, Johnson O, Meng QJ, Loudon ASI, Hastings MH. 2025. Quantitative measures of clock protein dynamics in the mouse suprachiasmatic nucleus extends the circadian time-keeping model. EMBO J Jul;44(13):3614–3644. doi: 10.1038/s44318-025-00426-z. Epub 2025 Apr 17. PMID: 40247113; PMCID: PMC12218236.

Strieder-Barboza C, Baker NA, Flesher CG, Karmakar M, Patel A, Lumeng CN, O’Rourke RW. 2020. Depot-specific adipocyte-extracellular matrix metabolic crosstalk in murine obesity. Adipocyte Dec;9(1):189–196. doi: 10.1080/21623945.2020.1749500. PMID: 32272860; PMCID: PMC7153651.

Sun Y, Rahbani JF, Jedrychowski MP, Riley CL, Vidoni S, Bogoslavski D, Hu B, Dumesic PA, Zeng X, Wang AB, et al. 2021. Mitochondrial TNAP controls thermogenesis by hydrolysis of phosphocreatine. Nature May;593(7860):580–585. doi: 10.1038/s41586-021-03533-z. Epub 2021 May 12. PMID: 33981039; PMCID: PMC8287965.

Takahashi JS. Transcriptional architecture of the mammalian circadian clock. 2017. Nat Rev Genet Mar;18(3):164–179. doi: 10.1038/nrg.2016.150. Epub 2016 Dec 19. PMID: 27990019; PMCID: PMC5501165.

The Gene Ontology Consortium; Aleksander SA, Balhoff J, Carbon S, Cherry JM, Drabkin HJ, Ebert D, Feuermann M, Gaudet P, Harris NL, Hill DP, et al. 2023. The Gene Ontology knowledgebase in 2023. Genetics May 4;224(1):iyad031. doi: 10.1093/genetics/iyad031. PMID: 36866529; PMCID: PMC10158837.

Thomas PD, Ebert D, Muruganujan A, Mushayahama T, Albou LP, Mi H. 2022. PANTHER: Making genome-scale phylogenetics accessible to all. Protein Sci Jan;31(1):8–22. doi:10.1002/pro.4218. Epub 2021 Nov 25. PMID: 34717010; PMCID: PMC8740835.

Thurley K, Herbst C, Wesener F, Koller B, Wallach T, Maier B, Kramer A, Westermark PO. 2017. Principles for circadian orchestration of metabolic pathways. Proc Natl Acad Sci U S A Feb 14;114(7):1572–1577. doi: 10.1073/pnas.1613103114. Epub 2017 Feb 3. PMID: 28159888; PMCID: PMC5321018.

Van Drunen R, Eckel-Mahan K. 2021. Circadian Rhythms of the Hypothalamus: From Function to Physiology. Clocks Sleep Feb 25;3(1):189–226. doi: 10.3390/clockssleep3010012. PMID:33668705; PMCID: PMC7931002.

van Rosmalen L, Deota S, Maier G, Le HD, Lin T, Ramasamy RK, Hut RA, Panda S. 2024. Energy balance drives diurnal and nocturnal brain transcriptome rhythms. Cell Rep Mar 26;43(3):113951. doi: 10.1016/j.celrep.2024.113951. Epub 2024 Mar 19. PMID: 38508192; PMCID: PMC11330649.

Vargas-Castillo A, Sun Y, Smythers AL, Grauvogel L, Dumesic PA, Emont MP, Tsai LT, Rosen ED, Zammit NW, Shaffer SM, et al. 2024. Development of a functional beige fat cell line uncovers independent subclasses of cells expressing UCP1 and the futile creatine cycle. Cell Metab Sep 3;36(9):2146–2155.e5. doi: 10.1016/j.cmet.2024.07.002. Epub 2024 Jul 30. PMID: 39084217; PMCID: PMC12005060.

Wu G, Anafi RC, Hughes ME, Kornacker K, Hogenesch JB. 2016. MetaCycle: an integrated R package to evaluate periodicity in large scale data. Bioinformatics Nov 1;32(21):3351–3353. doi: 10.1093/bioinformatics/btw405. Epub 2016 Jul 4. PMID: 27378304; PMCID: PMC5079475.

Xiong X, Lin Y, Lee J, Paul A, Yechoor V, Figueiro M, Ma K. 2021. Chronic circadian shift leads to adipose tissue inflammation and fibrosis. Mol Cell Endocrinol Feb 5;521:111110. doi:10.1016/j.mce.2020.111110. Epub 2020 Dec 4. PMID: 33285245; PMCID: PMC7799174.

Yang Loureiro Z, Solivan-Rivera J, Corvera S. 2022. Adipocyte Heterogeneity Underlying Adipose Tissue Functions. Endocrinology Jan 1;163(1):bqab138. doi: 10.1210/endocr/bqab138. PMID: 34223880; PMCID: PMC8660558.

Ye L, Wu J, Cohen P, Kazak L, Khandekar MJ, Jedrychowski MP, et al. 2013. Fat cells directly sense temperature to activate thermogenesis. Proc Natl Acad Sci U S A Jul 23;110(30):12480–12485. doi: 10.1073/pnas.1310261110. PMID: 23818608; PMCID: PMC3725076.

Zhang R, Lahens NF, Ballance HI, Hughes ME, Hogenesch JB. 2014. A circadian gene expression atlas in mammals: implications for biology and medicine. Proc Natl Acad Sci U S A Nov 11;111(45):16219–24. doi: 10.1073/pnas.1408886111. Epub 2014 Oct 27. PMID: 25349387; PMCID: PMC4234565.

Zhang R, Podtelezhnikov AA, Hogenesch JB, Anafi RC. 2016. Discovering Biology in Periodic Data through Phase Set Enrichment Analysis (PSEA). J Biol Rhythms Jun;31(3):244–57. doi: 10.1177/0748730416631895. Epub 2016 Mar 8. PMID: 26955841.

Zvonic S, Ptitsyn AA, Conrad SA, Scott LK, Floyd ZE, Kilroy G, Wu X, Goh BC, Mynatt RL, Gimble JM. 2006. Characterization of peripheral circadian clocks in adipose tissues. Diabetes Apr;55(4):962–70. doi: 10.2337/diabetes.55.04.06.db05-0873. PMID: 16567517.

