## Supplemental Figures for "Cell-Autonomous and Systemic Circadian Regulation of Gene Expression in Adipocytes"

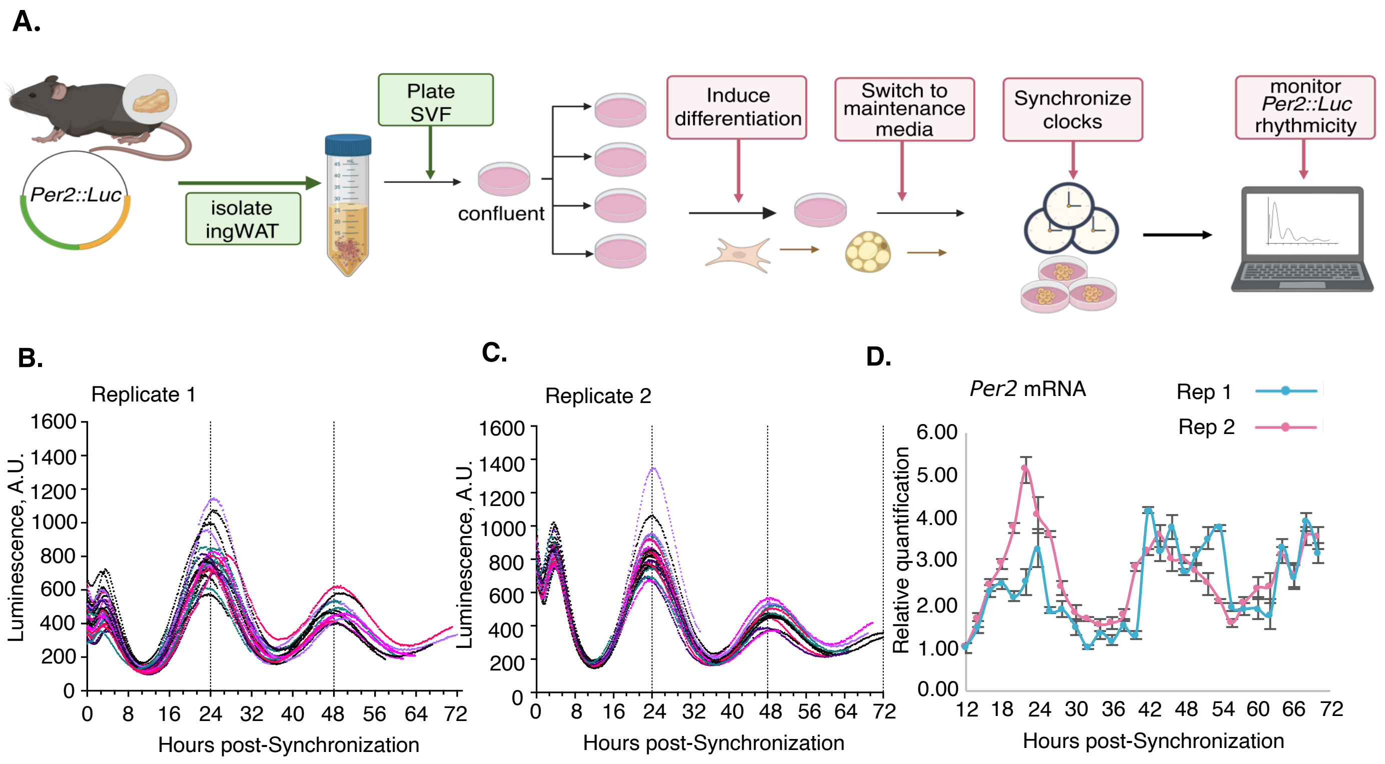


**Supplementary Fig. S1. IVDAs exhibit robust cell-specific *Per2* oscillations. (A)** Schematic of *Per2*::Luciferase reporter assay. Created in BioRender. Worthen, J. (2026) https://BioRender.com/uqxnwg2 **(B-C)** Luciferase traces for **(B)** replicate 1 and **(C)** replicate 2 with each line representing a culture dish which corresponds to one time point sample and each line terminates at the time when the sample was taken for RNA extraction. **(D)** qRT-PCR of *Per2* mRNA expression in Replicate 1 and 2 at 30 different time points over 60hrs following serum shock. Error bars depict standard error.

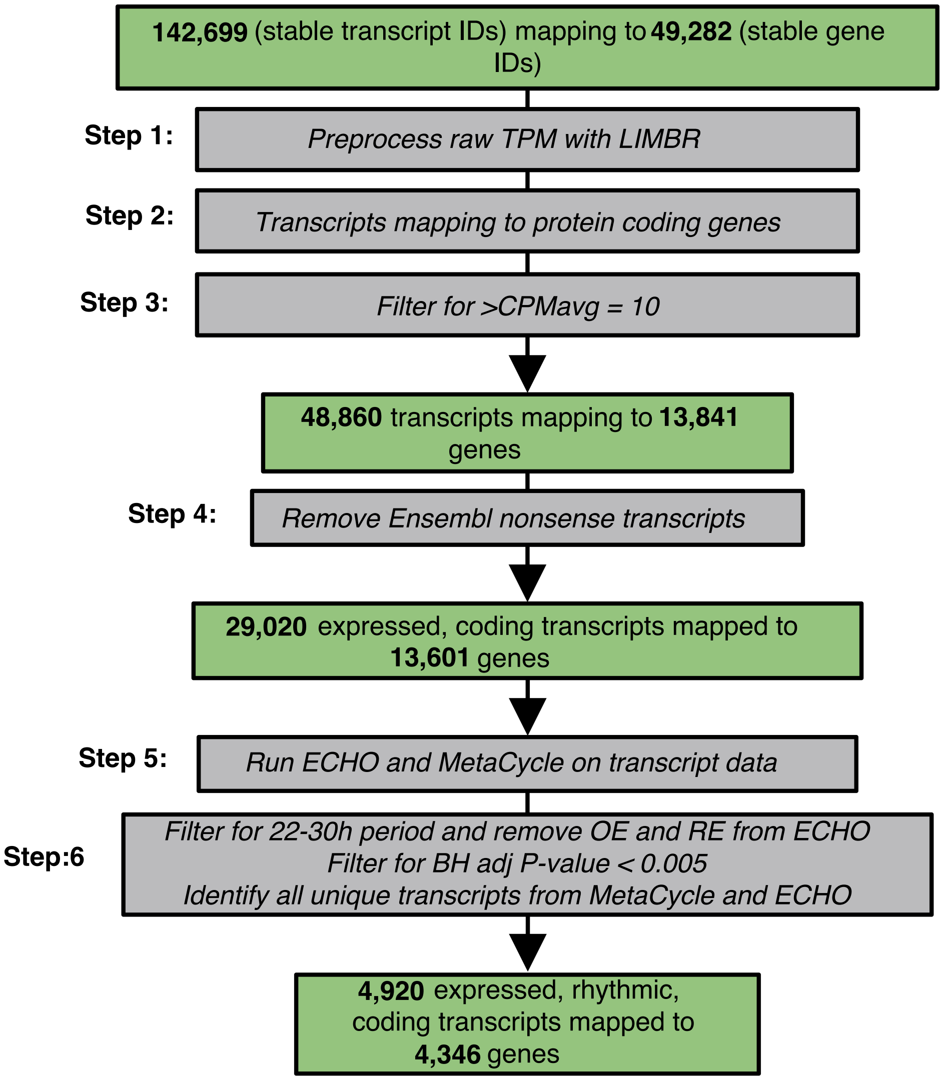


**Supplementary Fig. S2. Filtering strategy used to generate the circadian gene list for PSEA.** All 142,699 transcripts in the genome annotation, across all time points, were first preprocessed using LIMBR to remove batch effects. Transcripts were then mapped to protein-coding genes and filtered for average expression (CPMavg > 10) to remove lowly expressed transcripts. This cutoff was empirically determined based on olfactory gene expression (see Supplemental Methods). These steps yielded 48,860 transcripts mapping to 13,841 genes, which will be the focus of a separate manuscript. Next, transcripts were further filtered to retain only those predicted by Ensembl to produce a functional protein, thereby excluding nonsense and non-coding transcripts. This resulted in 29,020 transcripts mapping to 13,601 unique genes, which constitute the dataset analyzed in this manuscript. ECHO and MetaCycle were then applied to the preprocessed TPM data for these 29,020 transcripts. Rhythmic transcripts were filtered to retain periods between 22-30 h, with overexpressed (OE) and repressed (RE) oscillation types excluded. A significance threshold of BH-adjusted $P < 0.005$ was applied, yielding 4,920 rhythmic transcripts mapping to 4,346 genes. For genes producing multiple rhythmic transcripts, the most statistically significant transcript was selected. This final set of 4,346 genes and their associated peak phases (CT) was used as input for PSEA.

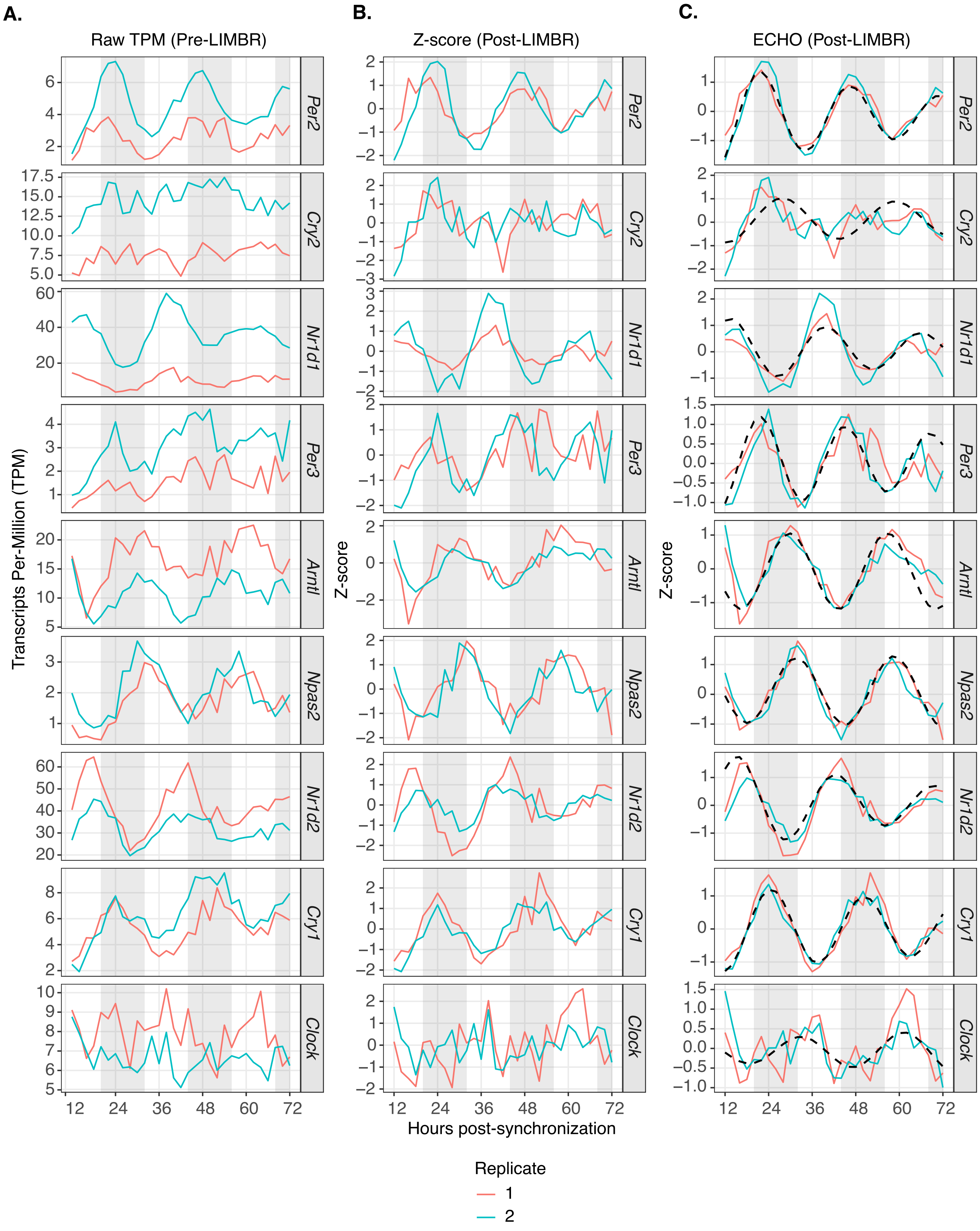


**Supplementary Fig. S3. Data processing and ECHO modeling of clock transcripts in IVDAs shows rhythmicity of clock genes**. RNA-seq expression data for clock transcripts presented as **(A)** Raw transcripts per million (TPM) values, **(B)** Z-scores, (C) Z-scores after pre-processing by LIMBR and **(D)** Modeled fits as determined by ECHO. Rep1 (pink) and Rep2 (blue) in IVDAs with the ECHO fitted curve shown in black dashed-line.

.

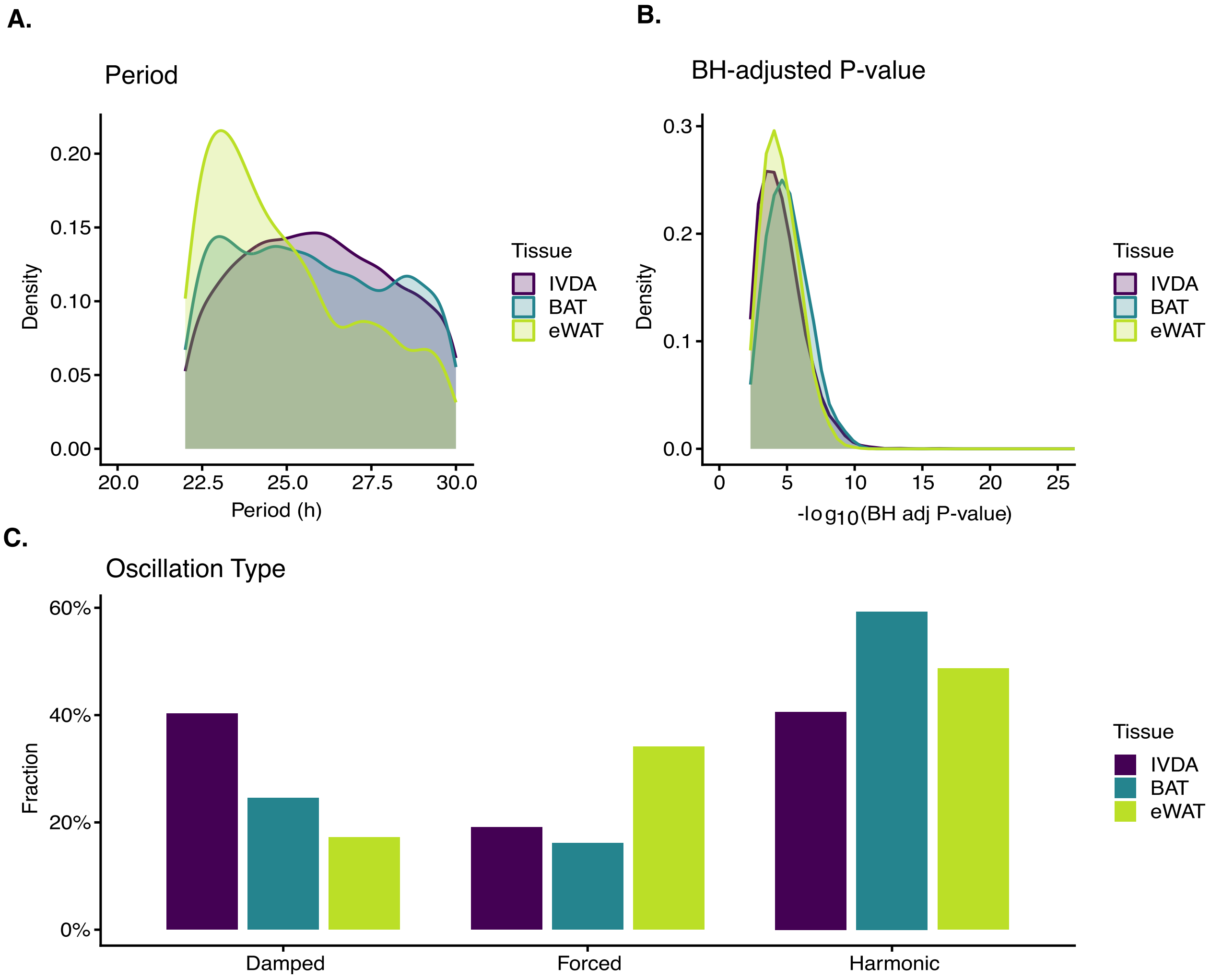


**Supplementary Fig. S4. Distribution of rhythmicity metrics across IVDAs, BAT and eWAT. (A-C)** Parameter density plots for IVDAs, BAT and eWAT (Zhang et al., 2014) depicting **(A)** Period distribution of transcripts oscillating within a 22-30h range and a BH adjusted p-value < 0.005, **(B)** BH adjusted p-value. Y-axes show density of the parameter being measured and x-axes show the unit of measurement for the parameter being shown, **(C)** Distribution of oscillation type by dataset**.**

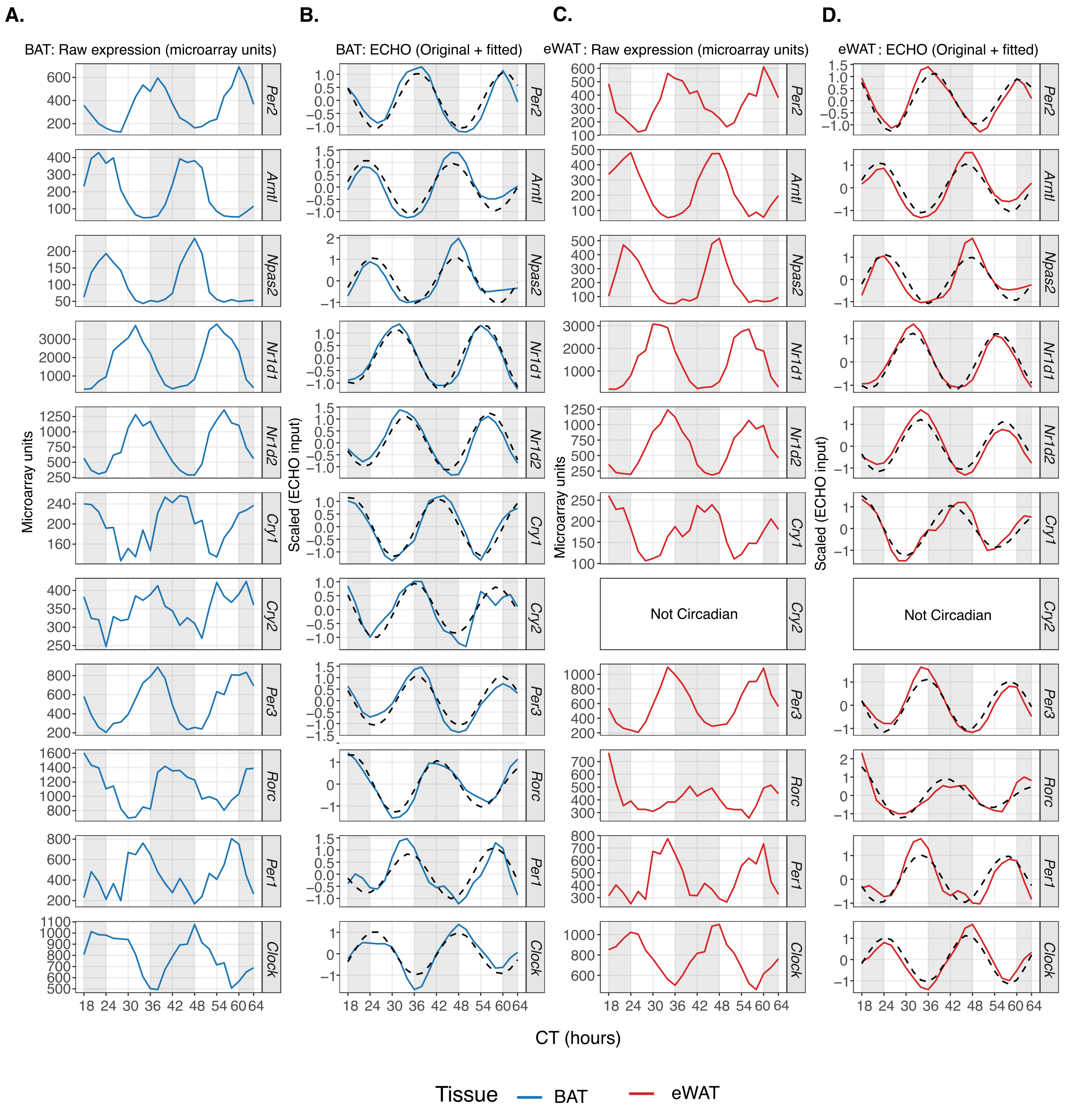


**Supplementary Fig. S5. Clock gene expression in BAT and eWAT (*in vivo*). (A,C)** Raw microarray expression profiles and **(B,D)** ECHO-modeled expression with fitted rhythmic curves for core circadian clock genes across circadian time (CT) in brown adipose tissue (BAT, blue) and white adipose tissue (eWAT, red). For ECHO panels, solid-colored lines show scaled expression values and black dashed lines indicate the ECHO-fitted rhythmic model. Genes are shown in the same order for BAT and eWAT. *Cry2* did not meet criteria for circadian rhythmicity in eWAT and is indicated as not circadian. Shaded regions denote the dark phase.

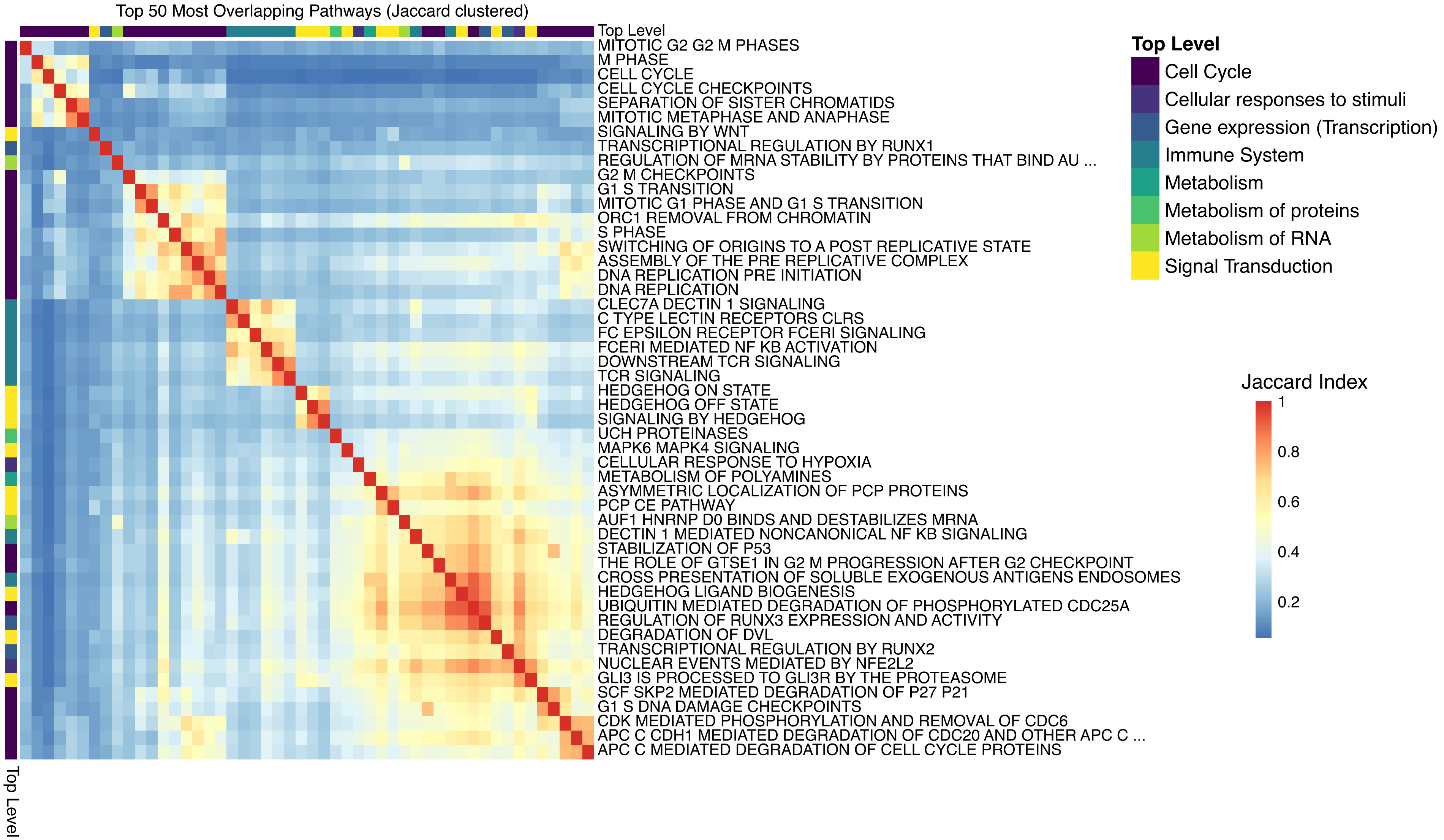


**Supplementary Fig. S6. PSEA leading-edge analysis shows that a common set of circadian genes drives enrichment only among cell-cycle related pathways.** Heatmap of pairwise Jaccard similarity (percentage of shared circadian genes) between the top 50 most overlapping phase-clustered pathways identified by PSEA in IVDA. Pathways are hierarchically clustered and colored by Reactome top-level category. A tight yellow cluster of high similarity corresponds to cell-cycle-related pathways, whereas most other categories show low overlap, indicating that phase clustering across functional groups in Fig. 2B arises from distinct sets of circadian genes rather than repeated detection of the same core drivers.

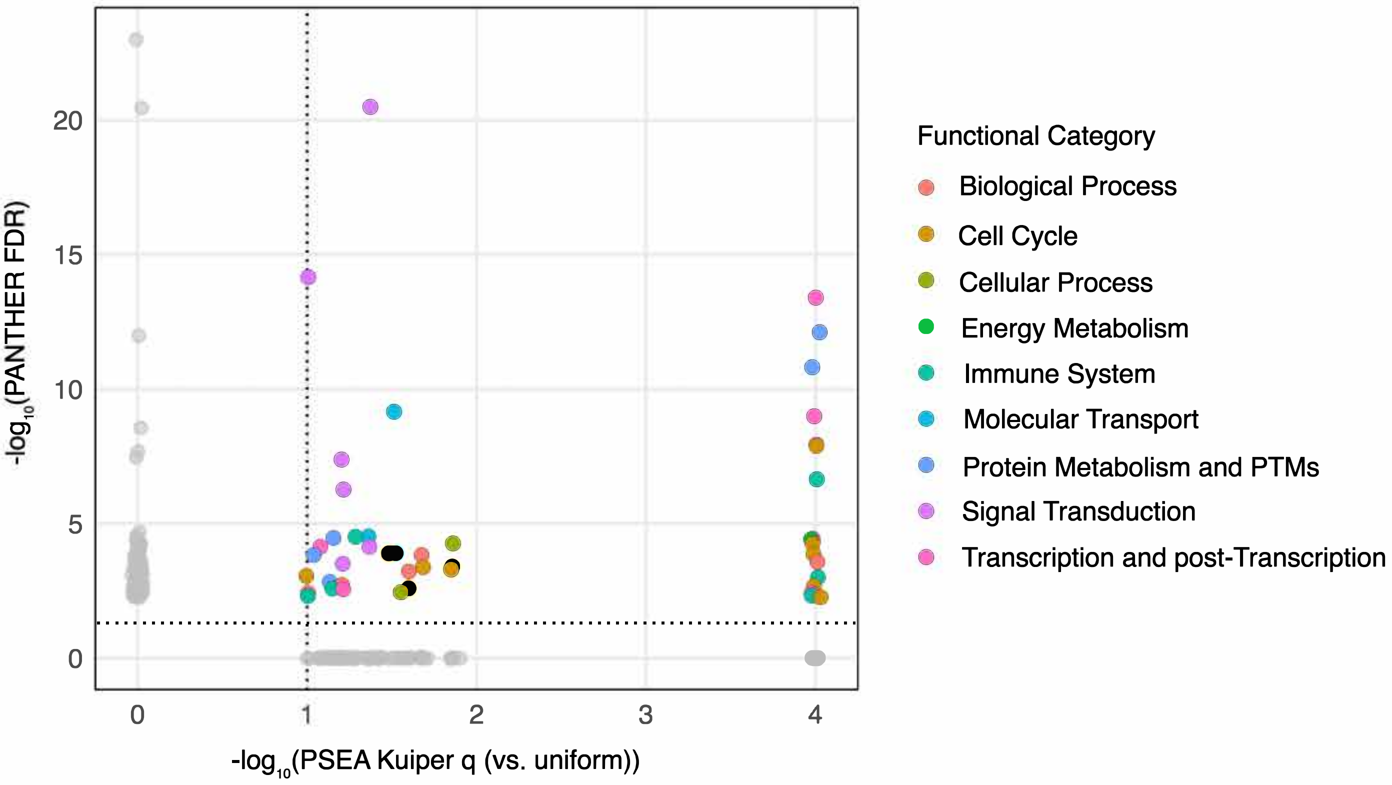


**Supplementary Fig. S7. Phase-clustered pathways detected by PSEA overlap with processes positively enriched for circadian genes detected by PANTHER in IVDAs**. Comparison of pathway significance between PSEA ($-\log_{10}$ Kuiper q, x-axis) and PANTHER ($-\log_{10}$ FDR, y-axis) for the pathways identified by each method. Each colored point represents a pathway that appears in both top-lists; color indicates its Reactome functional category. The dashed lines mark the respective significance thresholds (PSEA $q \leq 0.1$; PANTHER FDR $\leq 0.05$)**.**

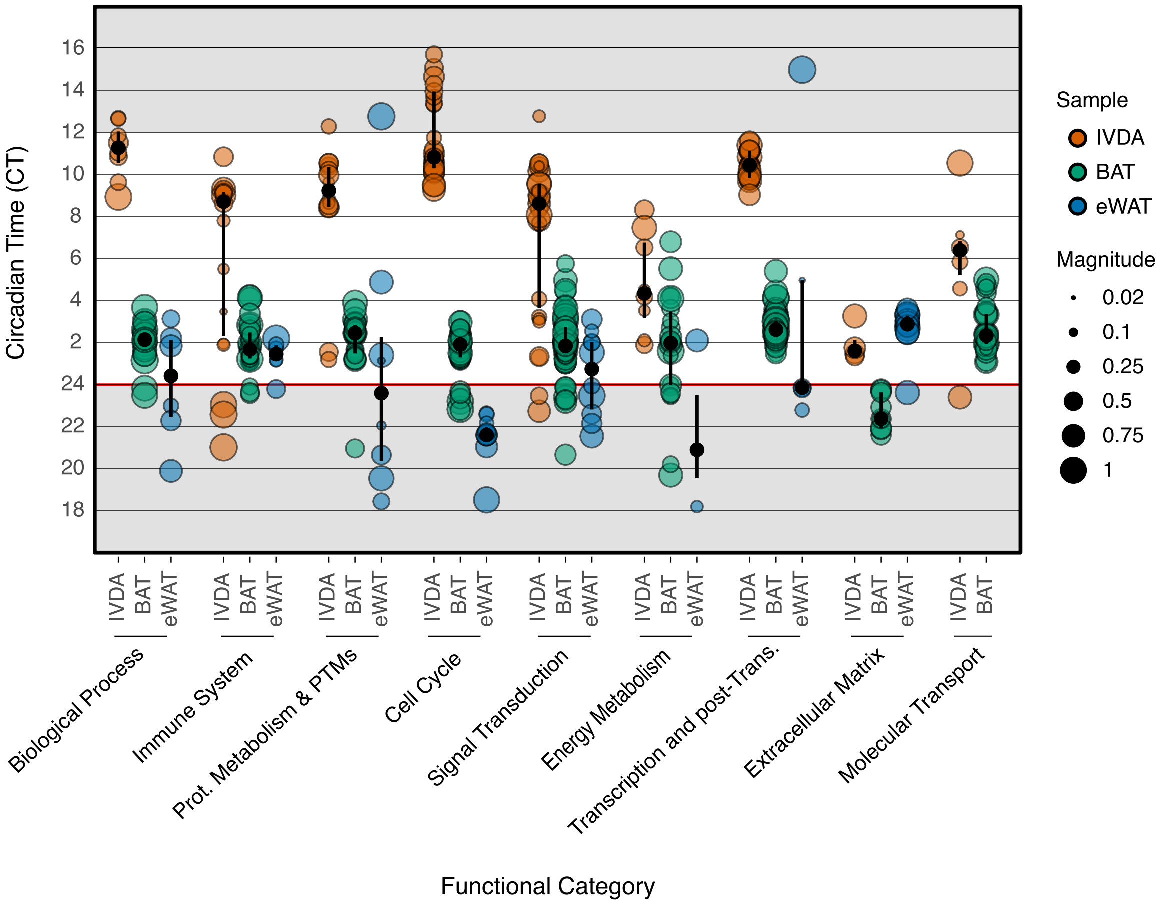


**Supplementary Fig. S8. Systemic cues contribute to the phase clustered magnitude of temporally regulated processes indicating they play a role in temporal cohesiveness**. The dot plot represents the phase clustered magnitude of circadianly regulated pathways for Reactome annotated gene sets determined to be temporally regulated by PSEA (Kuiper q -value < 0.1) in IVDAs, brown adipose tissue (BAT), and epididymal white adipose tissue (eWAT) (*in vivo* microarray data, Zhang et al., 2014) and partitioned into functional groups using the broader top-level Reactome hierarchal classifications for each pathway. The size of each point for IVDAs, BAT and eWAT represents the Kuiper q-value for that pathway. Statistics: The median is represented by the black dot, while Q1 and Q3 are denoted by the upper and lower black lines, respectively.

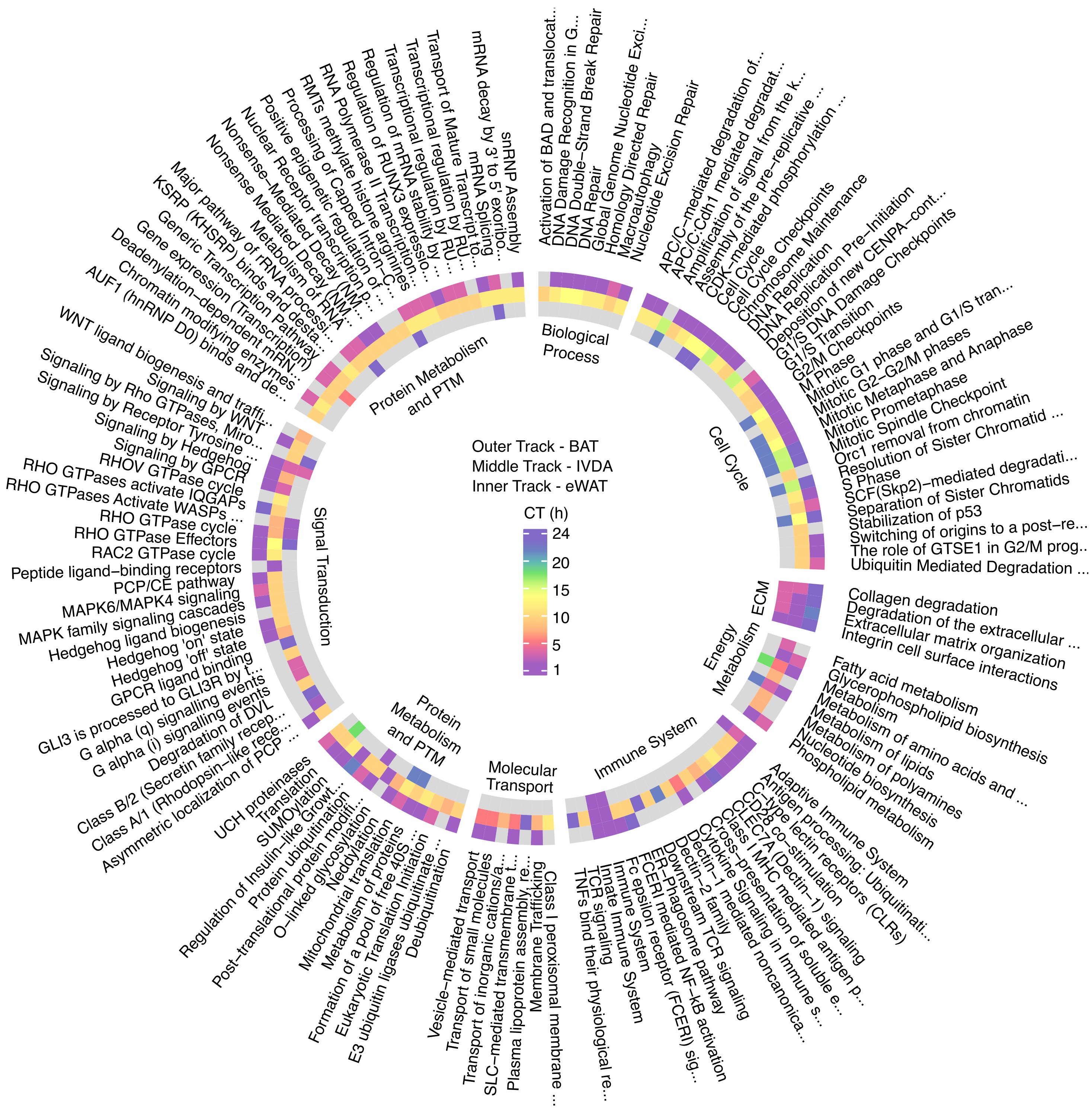


**Supplementary Fig. S9. Circular representation of phase-clustered Reactome pathways across adipose depots**. Each ring corresponds to one tissue (outer = BAT, middle = IVDA, inner = eWAT). Colored tiles indicate the circadian phase of each significantly rhythmic pathway (Kuiper $q < 0.1$) grouped by Reactome top-level category. Shared or overlapping pathways align radially, illustrating conserved versus tissue-specific timing across depots. The plot highlights conservation of circadian pathway identity, with phase shifts reflecting tissue- and cue-dependent temporal coordination *in vivo*.

.

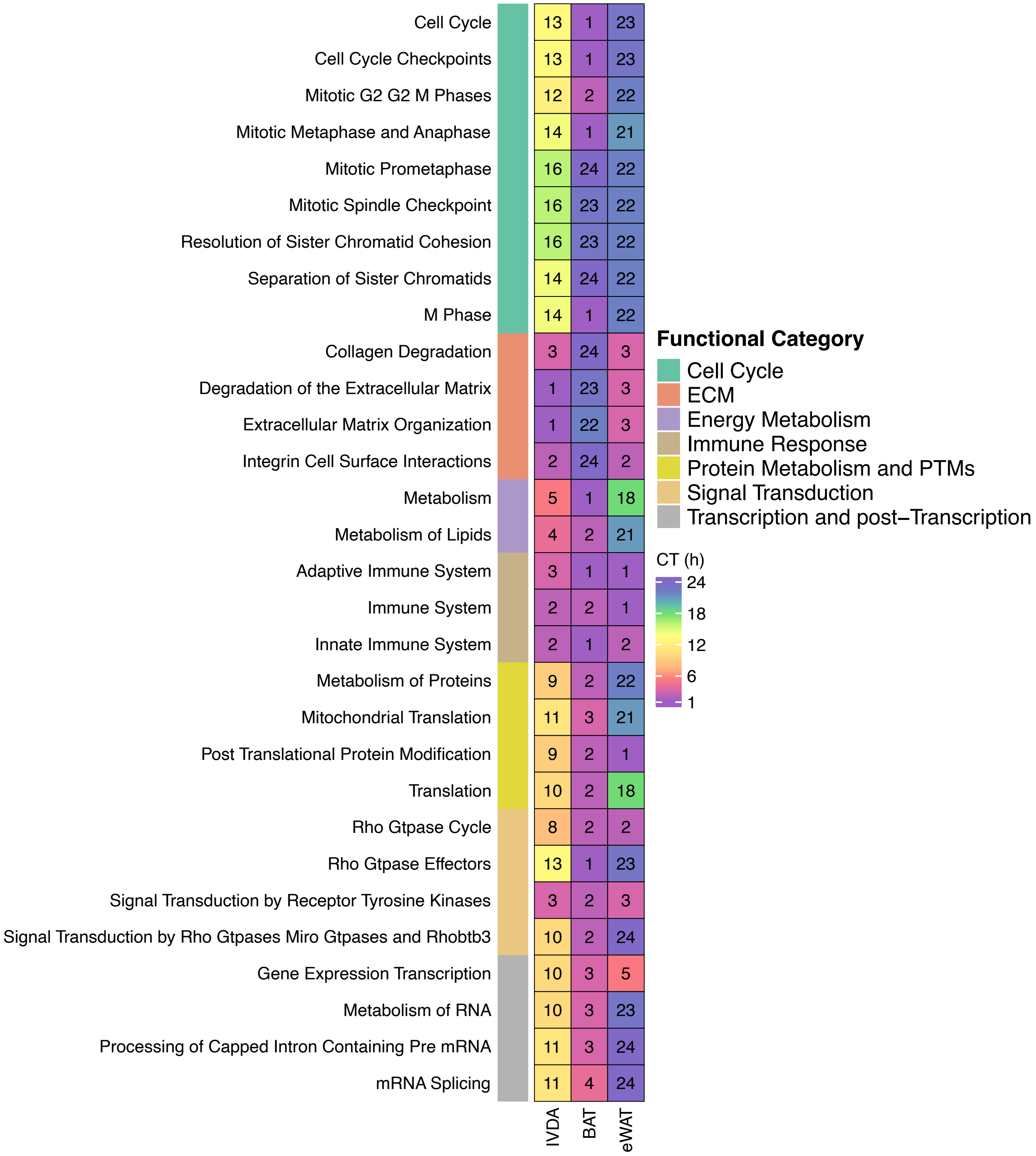


**Supplementary Fig. S10. Circadian peak times (CT) for the 30 Reactome pathways enriched in all three tissues**. Each row represents a Reactome pathway, and the three adjacent columns correspond to its peak circadian phase in IVDAs, BAT, and eWAT, respectively. Colors encode the circadian phase (1-24h). Pathways are organized by functional category (left color bar), including Cell Cycle, ECM, Energy Metabolism, Immune System, Protein Metabolism and PTMs, Signal Transduction, and Transcription and post-Transcription. Three pathways were excluded from the 33 overlap pathways in Fig. 2C because they did not fit into any functional category: Developmental Biology, Hemostasis, and Nervous System Development.

**Fig. S11A (Page 1)**

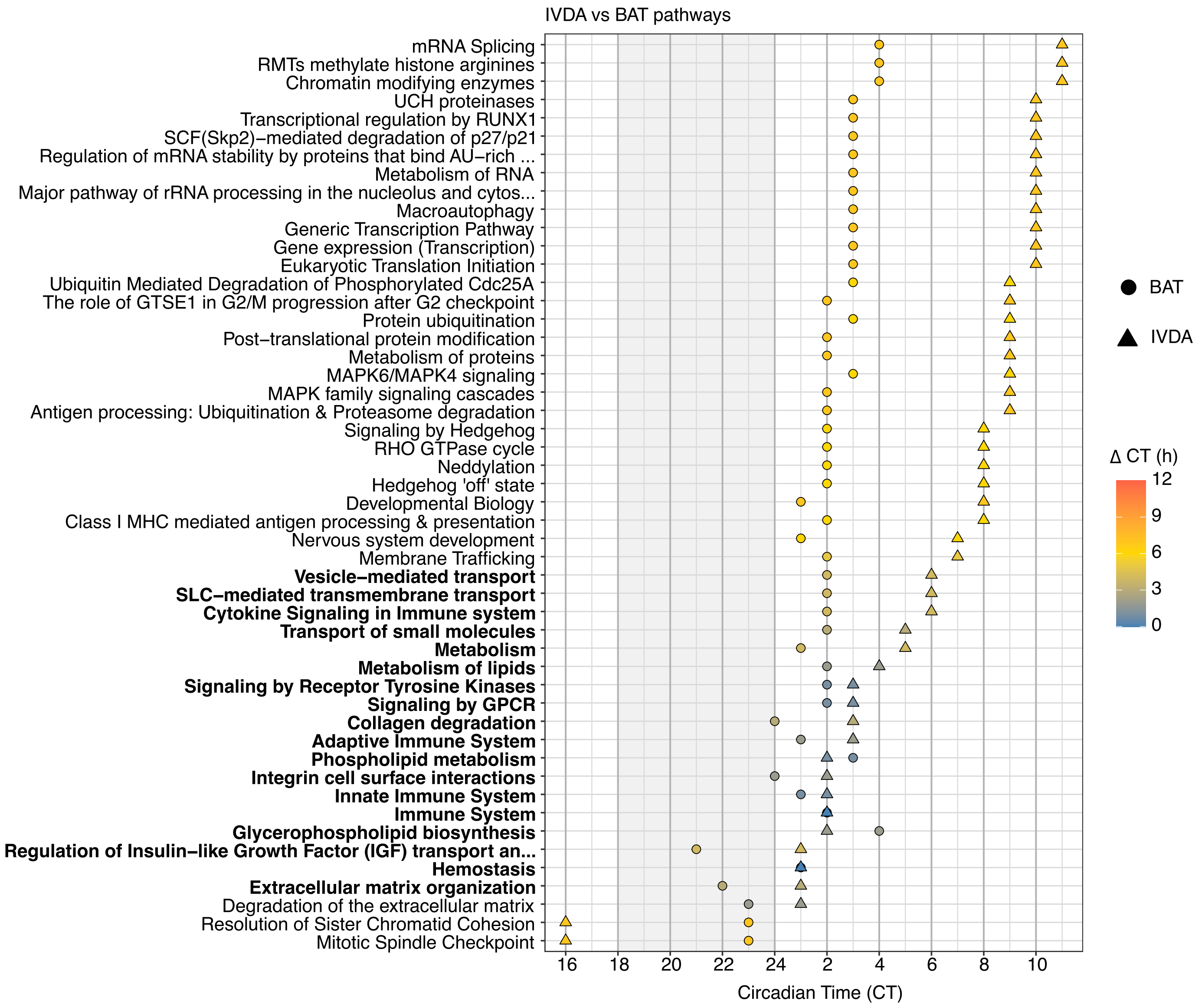

**Fig. S11A (Page 2)**

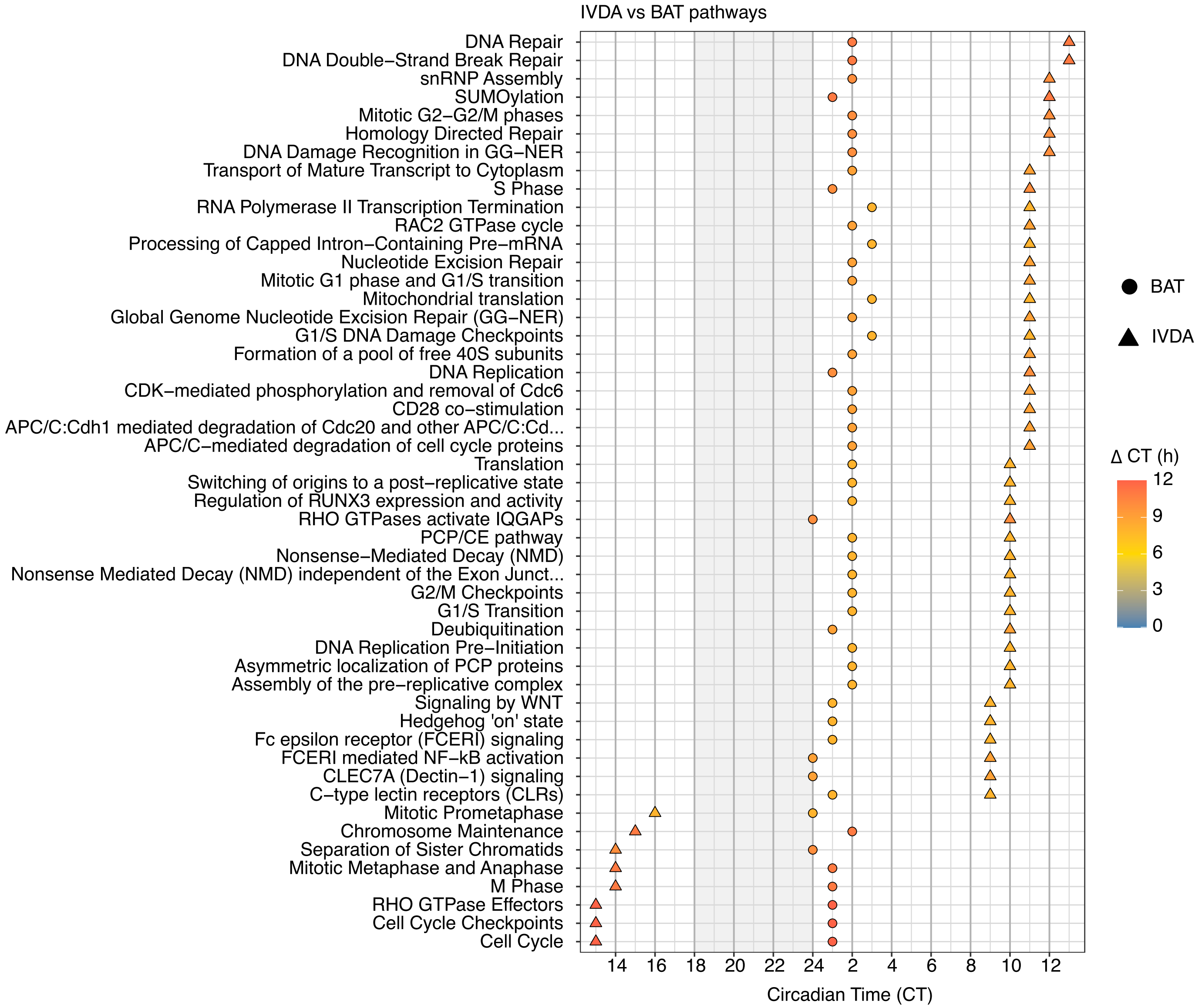

**Fig. S11B**

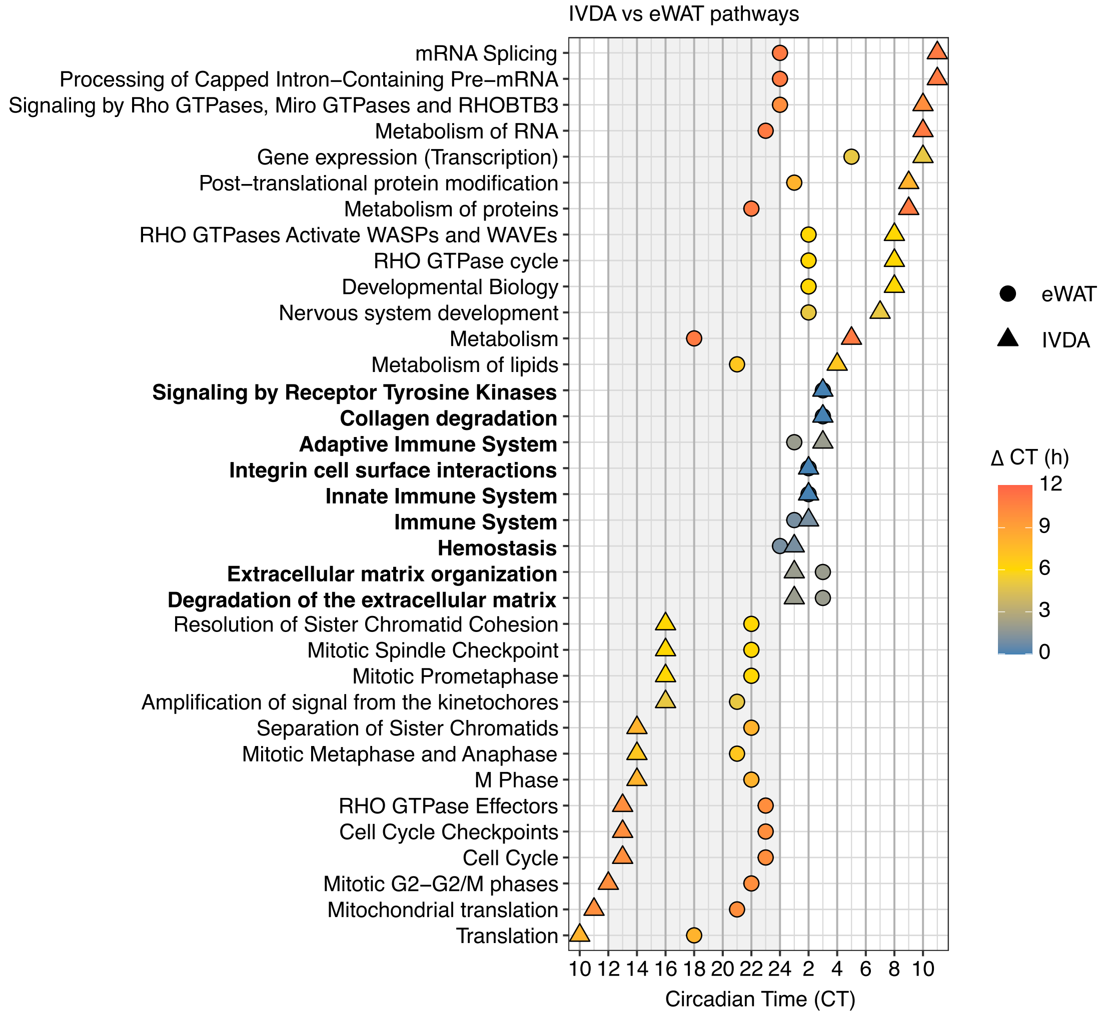


**Supplementary Fig. S11. Expanded comparison of phase-clustered pathways between IVDAs and adipose depots. (A)** Comparison of circadian timing (CT) for all overlapping phase-clustered pathways between IVDA and BAT, colored by the phase difference (ΔCT) from IVDA. **(B)** Same analysis for IVDA and eWAT. Each point represents the mean circadian time of peak pathway expression identified by PSEA (Kuiper q < 0.1). Circle and triangle symbols indicate IVDA and *in vivo* depots, respectively. Color scale denotes the magnitude of the phase shift (ΔCT = 0-12 h). Bolded pathways indicate they are in phase between the two datasets (ΔCT = 0-4 h). These pathways are also shown in Fig. 2D.

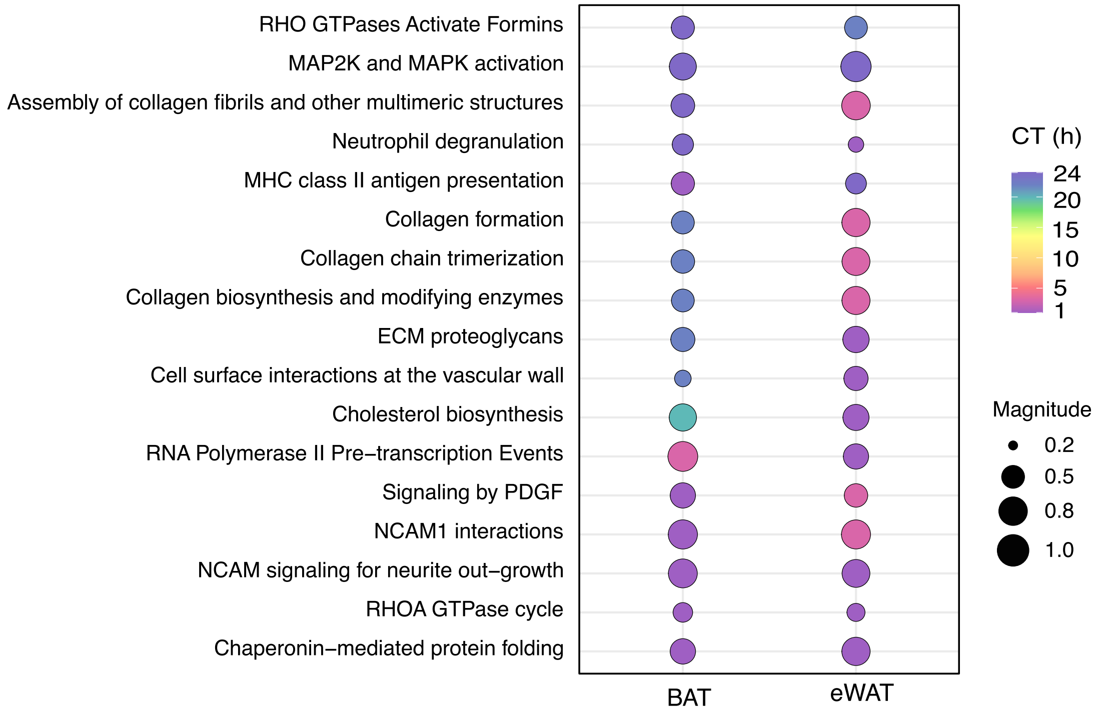


**Supplementary Fig. S12. Systemic circadian regulation of shared pathways in BAT and eWAT.** Phase dot plot of the shared BAT- eWAT pathways. Each row represents a pathway, and each column represents the source tissue (BAT, eWAT). The fill color corresponds to the circadian phase and size is represented by magnitude. Most shared pathways peak late-night early morning (CT20-4).

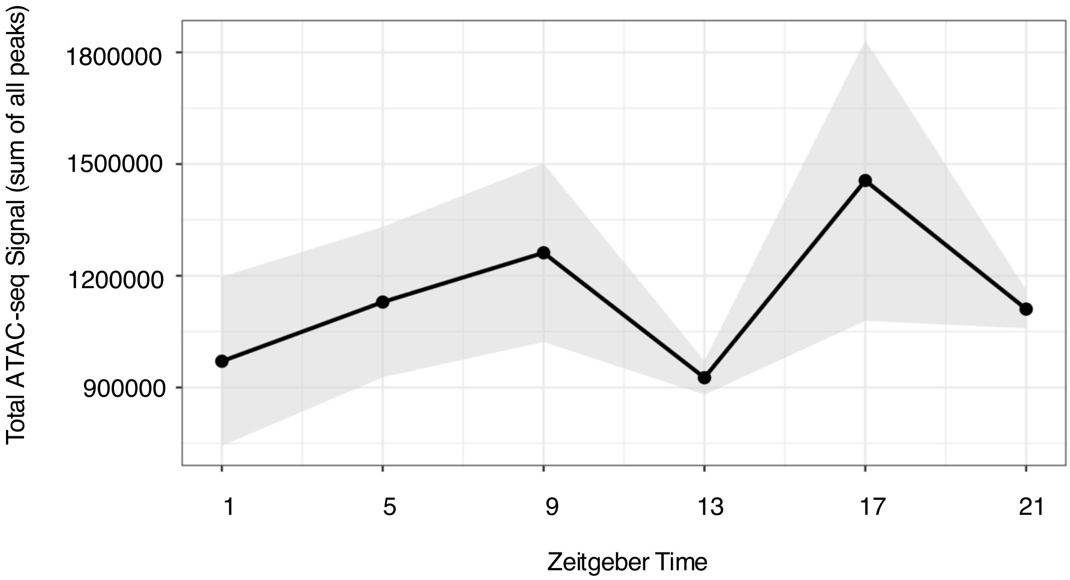


**Supplementary Fig. S13. Global chromatin accessibility varies across circadian time in BAT.** Genome-wide ATAC-seq signal in brown adipose tissue (BAT), calculated as the sum of normalized ATAC-seq peak intensities at each Zeitgeber time (ZT1, ZT5, ZT9, ZT13, ZT17, ZT21). Points represent the mean signal across biological replicates at each time point, with the black line connecting means to illustrate temporal trends. The shaded region denotes variability across replicates. These data indicate time of day dependent changes in overall chromatin accessibility in BAT.

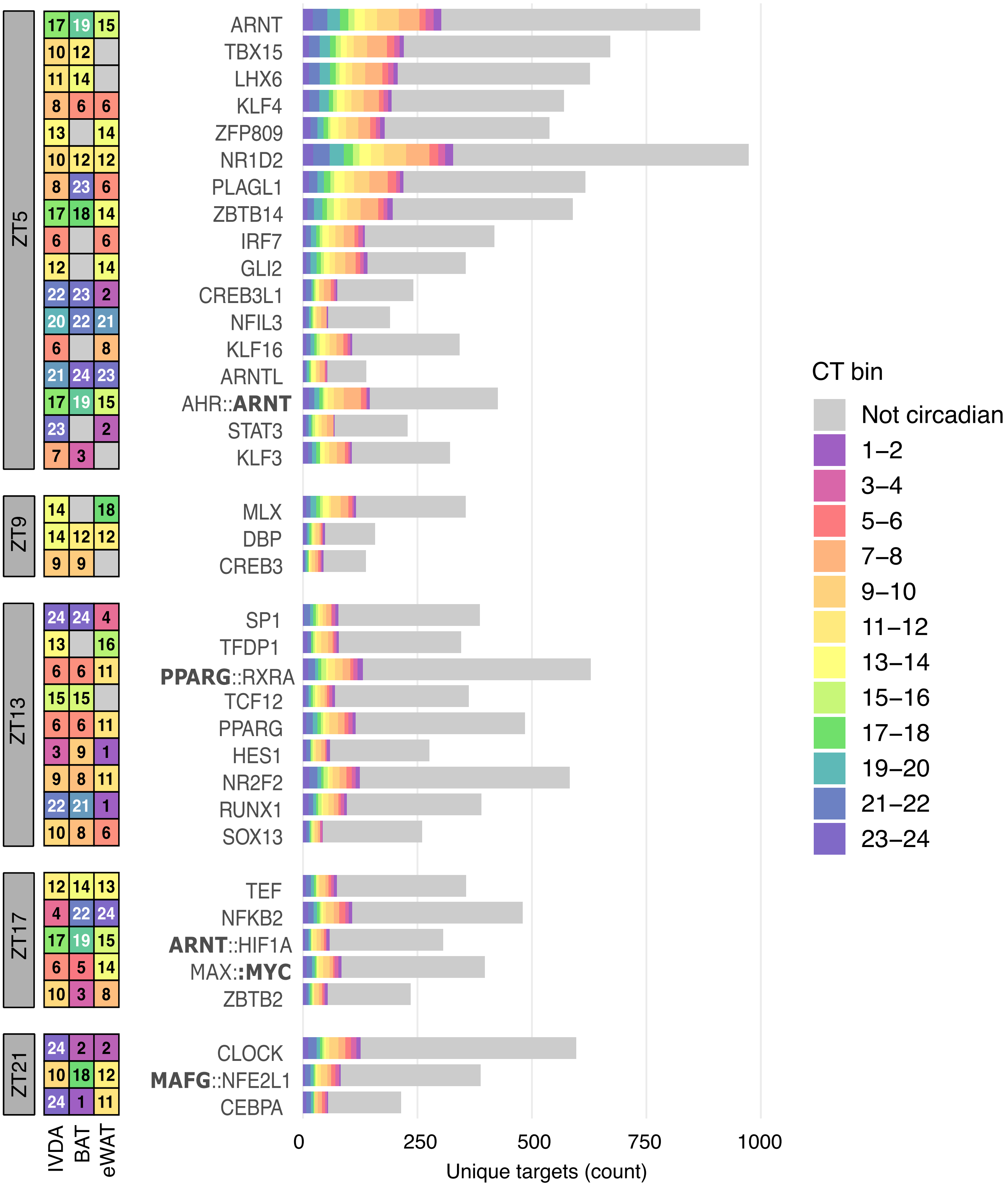


**Supplementary Fig. S14. Phase-aligned circadian transcription factor target gene distributions across adipose tissues**. The heatmap (left) is color-coded by the circadian time (CT) of peak activity in *in vitro* differentiated adipocytes (IVDAs), brown adipose tissue (BAT), and epididymal white adipose tissue (eWAT). Horizontal bars (right) show the number of unique predicted target genes for each transcription factor. Gray bars indicate targets that are not circadian in IVDAs. For transcription factor dimers, the CT value shown in the heatmap corresponds to the dimer partner indicated in bold. ZT, Zeitgeber time. No enrichment was found for ZT1.

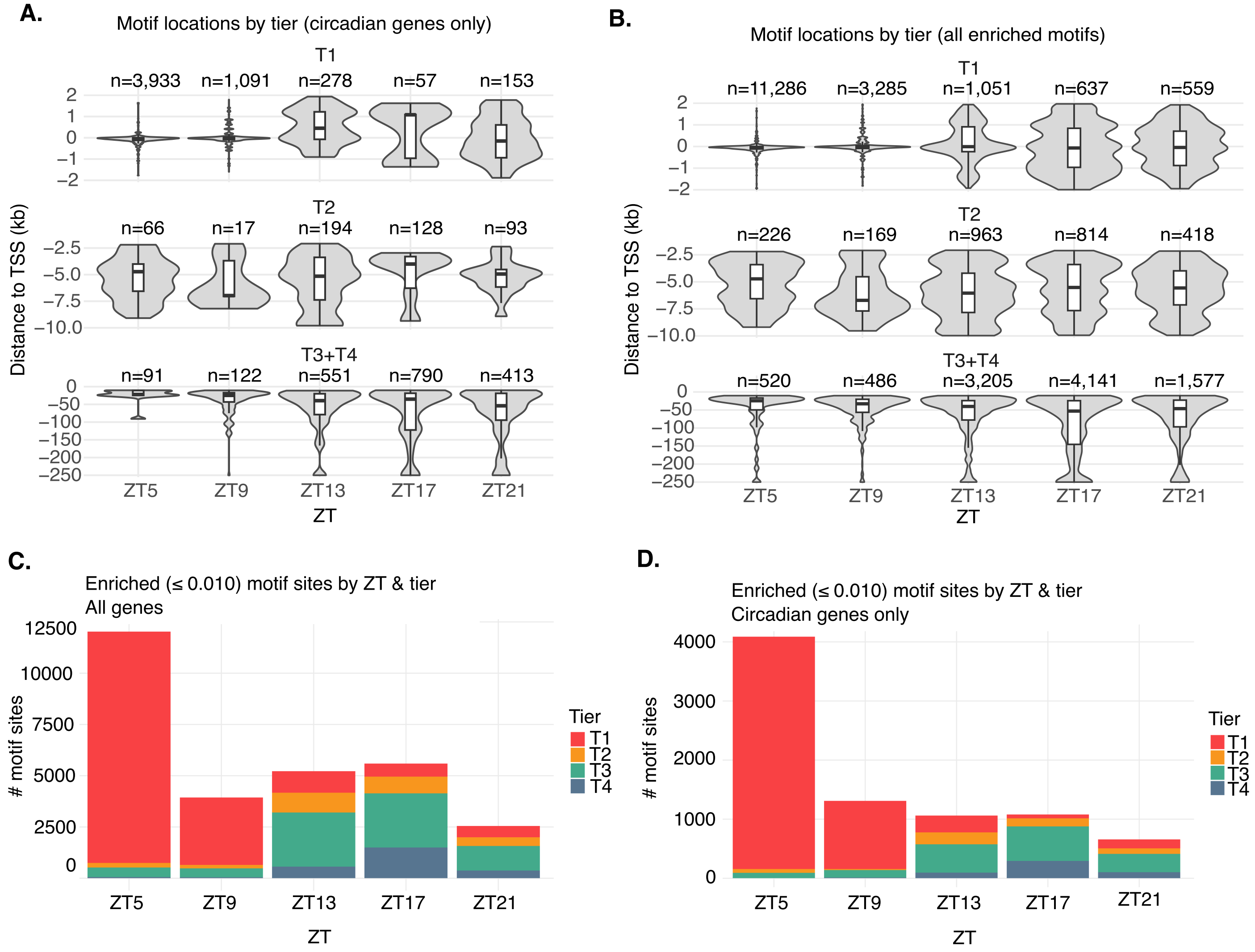


**Supplementary Fig. S15. Spatial dynamics of circadian TF motif enrichment relative to transcription start sites**. **(A-B)** Violin plots showing distance-to-TSS distribution for enriched motifs within the promoter (T1) and enhancer (T2, T3, T4) tiers of all genes (A) and circadian genes only **(B)** across ZT bins. **(C)** Total number of enriched (BH q ≤ 0.01) motif sites by circadian time and tier for all motifs. **(D)** Same as (C), restricted to circadian genes in IVDAs (ECHO + MetaCycle q ≤ 0.005). Tiers correspond to genomic distance from the nearest TSS: T1 (≤ 2 kb promoter), T2 (2-10 kb proximal), T3 (10-100 kb distal), T4 (> 100 kb far)**.**
