## Supplemental Methods for "Cell-Autonomous and Systemic Circadian Regulation of Gene Expression in Adipocytes"

**SUPPLEMENTARY METHODS**

**Animals and cell culture**

All procedures involving animals were performed according to protocols approved by the Institutional Animal Care and Use Committee (IACUC) of Dartmouth College. All mice were housed at 21 ± 2°C under a 12 h light/dark cycle and free access to normal chow (PicoLab Verified - 75 IF; 23.3% PRO, 12.9% FAT, 63.7% CHO) and water. Male mice homozygous for *Per2*::*Luc* aged 8 to 12 weeks (N=4) were used for all experiments. Originally a gift from the Takahashi Lab at Northwestern University, this line was rendered ~97% congenic with the strain C57BL/6J (Jackson Labs) through a SNP-based breeding program (Dartmouse) prior to being deposited at MMRRC as RRID:MMRRC_067372-MU. *Per2*::*Luc* is a "circadian reporter" line bearing a translational fusion of luciferase fused to the C-terminus of the clock protein PER2. All sacrifices were performed by forced $CO_2$ inhalation between 3-5 hours after lights on (lights on = 07:00 Eastern Standard Time). Lobes were transferred from a 1x PBS with penicillin-streptomycin to a collagenase solution prepared as follows - 0.1% BSA in 1x PBS without Mg+/Ca2+ rocked overnight in the cold to reduce bubbles. Then immediately before the inguinal stromal vascular fraction (SVF) from the bilateral inguinal white adipose lobes post-mortem excised within 20 minutes of death was added, Type I Collagenase (Sigma, Cat. # SR103, 1 mg/ml) and CalCl2 (Sigma, Cat. # 21115, 1 mM) were added to warmed BSA/PBS solution and filter-sterilized through a 0.22μm filter, ("collagenase solution"). Preadipocytes were isolated by mechanical dissociation with scissors for 4 minutes per every four lobes in collagenase digestion solution. Shaking incubation and centrifugation steps of SVF isolation were performed as in Yu et al. (2011) with minimal modifications. Cells were resuspended and seeded onto three 35 mm plastic-bottom tissue culture plates in freshly prepared “base media” containing DMEM/F-12 GlutaMAX (Gibco, Cat. # 10565-018) with 1% penicillin/streptomycin and 10% fetal bovine serum (Gibco, Lot # 2102306). Cells were incubated at 37°C in 5-6% $CO_2$ and passaged every 48 hours with a split ratio of approximately 1:2.8 for passage 1 and then 1:2 (32–34 dishes). In the final passage, cells were seeded into 35 mm plastic dishes (MatTek). Before confluence was reached, differentiation was induced with 1 μM synthetic glucocorticoid agonist dexamethasone (water-soluble, Sigma Cat. # D2915), 1 μM PPARg agonist rosiglitazone (Sigma Cat. # R2408 or Cayman Chemicals), 0.5 mM phosphodiesterase inhibitor IBMX (3-isobutyl-1-methylxanthine, Sigma Cat. # I5879); 33 μM biotin (Sigma Cat. # B4639), 100 nM insulin (from bovine pancreas, Sigma Cat. # I1882), 17 μM pantothenate (D-Pantothenic acid hemicalcium salt, Sigma Cat. # P5155) were also included in the media, "differentiation media". After 4 days of differentiation, dexamethasone and IBMX were removed from the media (now "maintenance media" comprised of rosiglitazone, insulin and B vitamins).

**Circadian bioluminescence monitoring and RNA-seq sample collection**

After four days in culture in maintenance media, *in vitro* differentiated adipocyte cells (IVDAs) were plump with lipid droplets by inspection with inverted light microscopy at 20x obj. IVDAs were then synchronized for 2-2.5 hours using “synchronization media”: 50% FBS, 50% "base media" and 10 μM forskolin (forskolin from *Coleus forskohlii*, Sigma Cat. # F3917). Synchronization was performed in batches of four dishes to minimize temperature changes. 0.01% bright d-luciferin (100 nM, GoldBio #LUCK-100), was added to “maintenance media” without dexamethasone (“recording maintenance media”) immediately before the final media switch, culture dishes were each topped with glass microscopy slides (Fisher Microscope Cover

Glass 220-38-99, 40 CIR-1) and sealed with Dow Corning high-vacuum silicone grease (Sigma Cat. # Z273554), then loaded into the luminometer (Lumicycle, Actimetrics, Evanston, IL). Changes in luciferase activity were monitored for four days with the Lumicycle placed in a 37°C, 0% $CO_2$ incubator (n=32 dishes) running LumiCycle version 1.4. Bioluminescence was continuously recorded for each sample for 1:11(min:sec) intervals, 6 intervals/recordings per hour with the following Lumicycle settings: Running average =1, smooth = 1, Butterworth order = 2, cutoff (hr) = 3 and 48. Data was subsequently analyzed with ClockLab. Sampling for two replicate time series began at 12 hours post-synchronization (HPS) to avoid artifacts immediately following serum shock (Balsalobre et al., 1998; Yagita et al., 2000) and continued every 2h with the final sample collected at 72 HPS. Period was calculated by averaging the trough-to-trough period of dishes with at least two peaks (n=18).

**RNA extraction and quantitative PCR**

Two independent time-courses were generated. Both replicates were sampled every 2 h over 60 h starting at 12 h post-synchronization. After washing the cells with PBS, TRIzol (1 ml; Thermo Fisher Scientific) was added, the cells were gently scraped, and cell lysates were collected in 1.5 ml RNase free tubes. Total RNA was extracted from cells using the Qiagen RNeasy Lipid Tissue Mini Kit (Cat. # 74804, Qiagen) according to manufacturer's protocol and RNA quality was assessed via fragment analyzer by the Dartmouth School of Medicine Genomics Shared Resource. cDNA was generated from 1μg of RNA by using the SuperScript™ IV First-Strand Synthesis System (Cat. # 18091050, ThermoFisher) and qPCR was performed on the CFX384 Touch Real-Time PCR Detection System (BioRad) using the iTaq Universal SYBR Green Supermix (Cat. # 725121, BioRad), according to the manufacturer's protocols. Relative expression quantification of the qPCR data was performed by using the ΔΔCT method. *Per2* was quantified to validate PER2 rhythmicity using *Eif2a* as the endogenous reference (Supplemental Figure 1D). Primers were obtained from Integrated DNA Technologies. Primer sequences are as follows: *Per2*_F: 5'- CTCCAGCGGAAACGAGAACTG-3'; *Per2*_R: 5' TTGGCAGACTGCTCACTACTG-3'; *Eif2a*_F: 5'-CAACGTGGCAGCCTTACA-3'; *Eif2a*_R: 5'- TTTCATGTCATAAAGTTGTAGGTTAGG-3'.

**RNA-sequencing and quantification**

Directional RNA-seq libraries were prepared using rRNA depletion. Libraries for the first replicate were sequenced on an Illumina NextSeq 500 platform with 100 bp paired-end reads (BGI group), while the second replicate was sequenced on an Illumina NovaSeq 6000 platform with 150 bp paired-end reads (Novogene Corporation) to a depth of at least 60M PE reads/sample. Reads mapping to any residual ribosomal rRNA were removed with SortMeRNA (version 4.3.4) (Kopylova et al., 2012) and reads were then mapped to the Mus musculus GRCm38 reference genome using the STAR (version 2.7) (Dobin et al., 2012) with the options --outSAMtype BAM SortedByCoordinate --outSAMunmapped Within --twopassMode Basic --outFilterMultimapNmax 1 --quantMode TranscriptomeSAM. The RNA-seq reads data were then quantified using RSEM version 1.3.1 (Bo and Dewey, 2011) with options --bam --paired-end --forward-prob 0 to produce counts for each and normalized to transcripts per million (TPM).

**Filtering of transcripts**

To reduce noise and exclude unexpressed and non-productive transcripts from our analysis, we filtered for genes with an average count per million (CPMavg) ≥ 10 and limited our analysis to those transcripts that map to protein-coding genes and removed nonsense transcripts from our analysis as these will be the focus of a future manuscript. To determine this noise cutoff, we measured the CPMavg of 3,001 transcripts mapping to the 1,164 olfactory receptor genes (GO:0004984) as these should not be expressed in adipose and 99.6% of these were below CPMavg=10. This filtering process yielded 29,020 transcripts mapping to 13,601 unique genes. For accurate comparison, we similarly filtered BAT and eWAT microarray data to remove transcriptional noise and include only expressed genes. We filtered for normalized expression >100 which was determined using olfactory gene expression as was done for our IVDAs; expression of the 1,164 olfactory receptor genes was ≤ 100 in ≤ 90% of BAT genes and ≤ 96% of eWAT genes. This resulted in a detected protein-coding transcriptome of 11,249 genes for BAT and 11,624 for eWAT. For both IVDAs and *in vivo* datasets, genes that did not meet these cutoffs were considered unexpressed and were excluded from our analysis.

**Preprocessing and rhythmicity tests**

IVDA expressions were preprocessed with LIMBR (Crowell et al., 2019) to remove batch effects and filtered to reduce noise and exclude unexpressed and non-productive transcripts as stated above. We then ran the Extended Circadian Harmonic Oscillator (ECHO) algorithm to identify rhythmic transcripts (De Los Santos et al., 2020) with freerun and all other default settings. ECHO has demonstrated superiority in its ability to detect steady-amplitude oscillations (harmonic) as well as forced and damped oscillations that respectively increase or decrease their amplitudes over time. As an addition measure of rhythmicity, we also used MetaCycle (Wu et al., 2016). Although the cells were demonstrably rhythmic based on real time *Per2::Luc* analysis, noise in transcript detection and limited frequency (only every 2 h) and duration (60 h) of sampling resulted in period estimates for known circadian genes (Figure 1D, Supplemental Figure 4) falling between 22 and 30 h. For this reason, genes were classified as rhythmic if one or more of their transcripts oscillated with a period of 22-30h and the Benjamini–Hochberg adjusted (BH adj) P-value (ECHO) or p-value (MetaCycle) was ≤ 0.005. This stringent cutoff ensures that only the most circadian transcripts were included in our analysis. For comparison, we similarly ran ECHO and MetaCycle on filtered microarray data for BAT and eWAT. We used the most circadian (highest significance) transcript from the union of ECHO and MetaCycle rhythmic transcripts to generate our list of CCGs on which we base our analysis. ECHO and MetaCycle results are provided in Supplemental Files 1-3.

We performed our analyses using transcript-level expression data rather than gene-level expression data because gene-level data frequently fails to accurately represent transcriptional activity (e.g., Yang et al., 2020 in Arabidopsis and Al-Athman et al., 2019 in baboon). In certain scenarios, although rhythmicity was detected at the gene-level, no circadian transcripts were detected for that gene. Conversely, there were cases where gene-level rhythmicity was absent, yet individual transcripts exhibited circadian expression. In the latter scenario, the absence of observed rhythmicity at the gene-level could be attributed to anti-phase circadian transcripts of equal abundance which “cancels out” the rhythmicity at the gene-level causing incorrect assignment of the gene as non-rhythmic. Consequently, we argue that investigating rhythmicity at the transcript-level takes advantage of the richness of the data set and yields a more precise estimate of how the circadian clock governs gene transcription, mitigating the compounding effects of transcription.

**Reactome pathway overrepresentation analysis**

Pathway overrepresentation analysis was performed on all circadian clock-controlled genes (CCGs) identified in the IVDA circadian transcriptome (Figures 1A, 2A, 3A, and 4A). Enrichment was assessed against a genome-wide background using Amigo/PANTHER with Reactome pathway gene set library (REACTOME v83, 2023). In addition, enrichment was performed for genes associated with rhythmic BAT chromatin accessibility, summarized across two major temporal clusters (Figure 6B) using the same library.

**PSEA to detect phase-clustered pathways**

We used phase set enrichment analysis (PSEA) (Zhang et al., 2016) to identify phase-clustered pathways using our final lists of circadian genes in IVDAs (4,346), BAT (4,061) and eWAT (3,157) as input (Figures 1A, 2A, 3A and 4A). We ran these against murine REACTOME gene set library from MSigDB (REACTOME v83, 2023). We also ran PSEA on only the IVDA CCGs in the REACTOME_METABOLISM pathway falling into either the subjective morning (CT5-16) or the subjective night (CT17-4) peaks (Figures 2C-D). We ran this list against a custom gene set file containing only pathways falling under the hierarchy of the REACTOME_METABOLISM top-level term. PSEA assigns a Kuiper score to each gene set; this metric quantifies the overall degree to which the average phase of expression of all CCGs in that gene set (e.g. CCGs involved in vesicle mediated transport) deviates from a uniform phase distribution. The Kuiper score, along with the vector-average-magnitude which reflects the temporal cohesiveness of the pathway to the assigned CT, serves as a measure of the overall rhythmic expression coherence within that gene set (Kuiper, 1962). We considered a pathway temporally regulated if its Kuiper q score was $\leq 0.1$ and items/set=5. A Kuiper q value $\leq 0.1$

corresponds to a slightly more stringent threshold than a Kuiper p value ≤ 0.05. Full PSEA results are provided as a Supplemental File 5.

**BAT ATAC-seq motif enrichment**

We searched nucleosome-free (accessible) chromatin regions from BAT ATAC-seq data published by Hepler et al. (2022) for enrichment of transcription factor (TF) binding motifs corresponding to circadian clock-controlled genes (CCGs) with DNA-binding activity (Gene Ontology term GO:0003070). This gene set was defined as factors that “interact selectively and non-covalently with a specific DNA sequence within the regulatory region of a gene in order to modulate transcription” and comprised 288 CCG-encoded TFs. TFs were classified as cell-autonomous if the phase difference between IVDAs and either *in vivo* adipose tissue dataset was less than 4 h, yielding 39 TFs. Processed data from the Hepler et al. study was retrieved from the GEO repository (GSE181443).

Position weight matrices (PWMs) for these TFs were obtained from the JASPAR database, restricted to high-confidence motifs with experimental support, resulting in 42 PWMs. No re-analysis of ATAC-seq rhythmicity was performed; peaks were considered rhythmic if they met the original publication’s criteria. BAT ATAC-seq data were generated at 4-h resolution across a 24-h cycle; to facilitate comparison with transcriptomic analyses, rhythmic ATAC-seq peaks were grouped into six 4-h Zeitgeber time (ZT) bins: ZT1 (ZT23-2), ZT5 (ZT3-6), ZT9 (ZT7-10), ZT13 (ZT11-14), ZT17 (ZT15-18), and ZT21 (ZT19-22).

Motif enrichment was assessed using motifmatchr (Schep, 2025), testing each ZT bin against all other circadian bins combined as background. Enrichment was evaluated for each TF motif using Fisher's exact test, followed by Benjamini-Hochberg correction for multiple testing. Motifs were considered significantly enriched at $p \leq 1 \times 10^{-3}$ and BH-adjusted $q \leq 0.01$.

### SUPPLEMENTARY METHODS REFERENCES


Ashburner M, Ball CA, Blake JA, Botstein D, Butler H, Cherry JM, Davis AP, Dolinski K, Dwight SS, Eppig JT, et al. 2000. Gene ontology: tool for the unification of biology. The Gene Ontology Consortium. *Nat Genet* May;25(1):25-9. doi: 10.1038/75556. PMID: 10802651; PMCID: PMC3037419.

Balsalobre A, Damiola F, Schibler U. 1998. A serum shock induces circadian gene expression in mammalian tissue culture cells.. *Cell* Jun 12;93(6):929-37. doi: 10.1016/s0092-8674(00)81199-x. PMID: 9635423.

Crowell AM, Greene CS, Loros JJ, Dunlap JC. 2019. Learning and Imputation for Mass-spec Bias Reduction (LIMBR). *Bioinformatics* May 1;35(9):1518-1526. doi: 10.1093/bioinformatics/bty828. PMID: 30247517; PMCID: PMC6499252.

De Los Santos H, Collins EJ, Mann C, Sagan AW, Jankowski MS, Bennett KP, Hurley JM. 2020. ECHO: an application for detection and analysis of oscillators identifies metabolic regulation on genome-wide circadian output. *Bioinformatics* Feb 1;36(3):773-781. doi: 10.1093/bioinformatics/btz617. PMID: 31384918; PMCID: PMC7523678.

Dobin A, Davis CA, Schlesinger F, Drenkow J, Zaleski C, Jha S, Batut P, Chaisson M, Gingeras TR. 2013. STAR: ultrafast universal RNA-seq aligner. *Bioinformatics* Jan 1;29(1):15-21. doi: 10.1093/bioinformatics/bts635. Epub 2012 Oct 25. PMID: 23104886; PMCID: PMC3530905.

El-Athman R, Knezevic D, Fuhr L, Relógio A. 2019. A Computational Analysis of Alternative Splicing across Mammalian Tissues Reveals Circadian and Ultradian Rhythms in Splicing Events. *Int J Mol Sci* Aug 15;20(16):3977. doi: 10.3390/ijms20163977. PMID: 31443305; PMCID: PMC6721216.

Hepler C, Weidemann BJ, Waldeck NJ, Marcheva B, Cedernaes J, Thorne AK, Kobayashi Y, Nozawa R, Newman MV, Gao P, Shao M, Ramsey KM, Gupta RK, Bass J. 2022. Time-restricted feeding mitigates obesity through adipocyte thermogenesis. *Science* Oct 21;378(6617):276-284. doi: 10.1126/science.abl8007. Epub 2022 Oct 20. PMID: 36264811; PMCID: PMC10150371.

Kopylova E, Noé L, Touzet H. 2012. SortMeRNA: fast and accurate filtering of ribosomal RNAs in metatranscriptomic data. *Bioinformatics* Dec 15;28(24):3211–3217. doi:10.1093/bioinformatics/bts611. PMID: 23071270

Kuiper NH. 1960. Tests concerning random points on a circle. *Proc Koninklijke Nederlandse Akademie van Wetenschappen Ser A* 63:38–47. doi: 10.1016/S1385-7258(60)50006-0.

Li B, Dewey CN. 2011. RSEM: accurate transcript quantification from RNA-Seq data with or without a reference genome. *BMC Bioinformatics* Aug 4;12:323. doi: 10.1186/1471-2105-12-323. PMID: 21816040; PMCID: PMC3163565.

Rauluseviciute I, Riudavets-Puig R, Blanc-Mathieu R, Castro-Mondragon JA, Ferenc K, Kumar V, Lemma RB, Lucas J, Chèneby J, Baranasic D, et al. 2024. JASPAR 2024: 20th anniversary of the open-access database of transcription factor binding profiles. *Nucleic Acids* Res Jan 5;52(D1):D174–D182. PMID: 38090905; PMCID: PMC10767964.

Schep A. 2025. motifmatchr: fast motif matching in R. *Bioconductor* R package version 1.30.0. doi:10.18129/B9.bioc.motifmatchr. Available at: https://bioconductor.org/packages/motifmatchr

Subramanian A, Tamayo P, Mootha VK, Mukherjee S, Ebert BL, Gillette MA, Paulovich A, Pomeroy SL, Golub TR, Lander ES, et al. 2005. Gene set enrichment analysis: a knowledge-based approach for interpreting genome-wide expression profiles. *Proc Natl Acad Sci U S A* Oct 25;102(43):15545-50. doi: 10.1073/pnas.0506580102. Epub 2005 Sep 30. PMID: 16199517; PMCID: PMC1239896.

Wu G, Anafi RC, Hughes ME, Kornacker K, Hogenesch JB. 2016. MetaCycle: an integrated R package to evaluate periodicity in large scale data. *Bioinformatics* Nov;32(21):3351–3353. doi:10.1093/bioinformatics/btw405. PMID: 27378304; PMCID: PMC5079475.

Yagita K, Okamura H. 2000. Forskolin induces circadian gene expression of rPer1, rPer2 and dbp in mammalian rat-1 fibroblasts. *FEBS Lett* Jan 7;465(1):79-82. doi: 10.1016/s0014-5793(99)01724-x. PMID: 10620710.

Yang Y, Li Y, Sancar A, Oztas O. 2020. The circadian clock shapes the Arabidopsis transcriptome by regulating alternative splicing and alternative polyadenylation. *J Biol Chem* May 29;295(22):7608-7619. doi: 10.1074/jbc.RA120.013513. Epub 2020 Apr 17. PMID: 32303634; PMCID: PMC7261790.

Yu G, Wu X, Kilroy G, Halvorsen YD, Gimble JM, Floyd ZE. 2011. Isolation of murine adipose-derived stem cells. *Methods Mol Biol* 2011;702:29-36. doi: 10.1007/978-1-61737-960-4_3. PMID: 21082392.

Zhang R, Podtelezhnikov AA, Hogenesch JB, Anafi RC. 2016. Discovering Biology in Periodic Data through Phase Set Enrichment Analysis (PSEA). *J Biol Rhythms* Jun;31(3):244-57. doi: 10.1177/0748730416631895. Epub 2016 Mar 8. PMID: 26955841.
