## Supplemental Table 1 for "Cell-Autonomous and Systemic Circadian Regulation of Gene Expression in Adipocytes"

| **BH adj. p-value** | | **Total circadian genes** | **Circadian BMAL1 targets** | **Total % of circadian genes** |
| --- | --- | --- | --- | --- |
| *≤ 0.05* | 6,030 | | 2,203 | 36.5% |
| *≤ 0.01* | 5,282 | | 1,997 | 37.8% |
| *≤ 0.005* | 4,347 | | 1,961 | 45.1% |
| *≤ 0.001* | 3,651 | | 1,679 | 46.0% |
| *≤ 0.0001* | 2,550 | | 1,190 | 46.7% |

**Supplemental Table 1. Percentage of BMAL1 target genes in CCGs at different significance cutoffs.** Shows the percentage of BMAL1 target genes found in lists of CCGs meeting the BH *p-value* cutoffs of ≤ 0.05, ≤ 0.01, ≤ 0.005, ≤ 0.001, ≤ 0.00001. List of BMAL1 target genes was obtained from Hepler et al., 2022 BMAL1 ChIP-seq in inguinal white adipose tissue (iWAT) filtered for those genes with BMAL1 peaks located -10kb to 1kb from their transcriptional start site.
