## Supplemental Table 2 for "Cell-Autonomous and Systemic Circadian Regulation of Gene Expression in Adipocytes"

| **Gene** | **IVDAs** | **BAT** | **eWAT** |
| --- | --- | --- | --- |
| ***Arntl*** | 21 | 24 | 23 |
| ***Clock*** | 24 | 2 | 2 |
| ***Cry1*** | 17 | 19 | 17 |
| ***Cry2*** | NA | 16 | NA |
| ***Npas2*** | 23 | 2 | 1 |
| ***Nr1d1*** | 7 | 8 | 8 |
| ***Nr1d2*** | 10 | 12 | 12 |
| ***Per1*** | NA | 12 | 12 |
| ***Per2*** | 14 | 14 | 14 |
| ***Per3*** | 14 | 15 | 14 |
| ***Rora*** | NA | NA | 15 |
| ***Rorc*** | NA | 20 | 17 |

**Supplemental Table 2. Clocks of IVDAs are in-phase with that of *in vivo* BAT and eWAT.** Table shows the phase in circadian time (CT) of clock transcripts from IVDAs, brown adipose tissue (BAT) and epididymal white adipose tissue (eWAT) (*in vivo* microarray data, Zhang et al., 2014). N/A indicates there are no detected rhythmic transcripts for that gene.
